# Expected Distractor Context Biases the Attentional Template for Target Shapes

**DOI:** 10.1101/2022.10.18.512686

**Authors:** Maëlle Lerebourg, Floris P. de Lange, Marius V. Peelen

## Abstract

Visual search is supported by an internal representation of the target, the attentional template. However, which features are diagnostic of target presence critically depends on the distractors. Accordingly, previous research showed that consistent distractor context shapes the attentional template for simple targets, with the template emphasizing diagnostic dimensions (e.g., colour or orientation) in blocks of trials. Here, we investigated how distractor expectations bias attentional templates for complex shapes, and tested whether such biases reflect inter-trial priming or can be instantiated flexibly. Participants searched for novel shapes (cued by name) in two probabilistic distractor contexts: either the target’s orientation or rectilinearity was unique (80% validity). Across four experiments, performance was better when the distractor context was expected, indicating that target features in the expected diagnostic dimension were emphasized. Attentional templates were biased by distractor expectations when distractor context was blocked, also for participants reporting no awareness of the manipulation. Interestingly, attentional templates were also biased when distractor context was cued on a trial-by-trial basis, but only when the two contexts were consistently presented at distinct spatial locations. These results show that attentional templates can flexibly and adaptively incorporate expectations about target-distractor relations when looking for the same object in different contexts.

**Public significance statement:** When searching for an object (e.g., a green ball), the visual features that distinguish it from distractor objects depend on the features of these distractors (e.g., when searching among plants, its green colour is not useful to find the target). Here, we asked participants to search for novel shapes in contexts where different dimensions of the shapes were unique. We show that people learn which features are diagnostic in these distractor contexts and flexibly use expectations about the features that are diagnostic of the target to efficiently guide search.

## Introduction

Goal-directed behaviour often requires visually searching for relevant objects in our environment. To find such target objects, we need to know the visual features characterizing them. For example, when searching for one’s favourite green coffee mug among many other objects, we keep in mind the mug’s specific colour and shape. The internal representation of the target is often called the *attentional template* or *search template* (Battistoni et al., 2017; Duncan & Humphreys, 1989; Eimer, 2014; Wolfe, 2021): a working memory representation of visual features that characterize the target. Attentional templates may act as a top-down bias (Desimone & Duncan, 1995), supporting search by guiding spatial attention to the target and/or supporting target identification (Hout & Goldinger, 2014; Wolfe, 2021).

To allow for efficient search, attentional templates should ideally include those features that distinguish the target from potential distractor objects (Duncan & Humphreys, 1989; Geng & Witkowski, 2019; Navalpakkam & Itti, 2007). Which features are diagnostic depends not only on the target itself but also on the current distractors: while looking for green objects may help to quickly find the green coffee mug on one’s office desk, such a template would become much less useful if the mug was placed on the windowsill next to multiple green plants. Thus, when the distractor context is known (or predictable), this calls for a template that is shaped by expected distractors, flexibly highlighting diagnostic target features depending on context.

This idea has now gained some support. Generally, the statistical distribution of distractor features can be accurately learnt during search (Chetverikov et al., 2016, 2017). Based on this distractor knowledge, the template may be modulated accordingly, as evidenced by a range of studies investigating how attentional capture or memory representations of the target are influenced by predictable distractors. Within one specific feature dimension, such as colour or orientation, the attentional template could be shifted away from distractors (e.g., not the mug’s particular shade of green but a slightly darker green, compared to all plants on the windowsill, Geng et al., 2017; Kerzel, 2020; Scolari et al., 2012; Scolari & Serences, 2009; Yu & Geng, 2019) or defined entirely relative to the distractors, without taking any specific feature value (e.g., a template for ‘greener’, but not a specific shade of green) (S. I. Becker, 2010b, 2010b; S. I. Becker et al., 2013). These studies therefore provide evidence that attentional templates can be influenced by distractor context, allowing observers to distinguish targets from distractors more efficiently when these are characterized only by small feature differences. Context may affect processing at different search stages, and while there is evidence for context effects on attentional selection (S. I. Becker et al., 2010; Kerzel, 2020; Scolari & Serences, 2009) and sensory processing (Scolari & Serences, 2009), some of these effects could also reflect changes in later target identification and decision processes (Hamblin-Frohman & Becker, 2021; Yu et al., 2022).

Most targets and distractors are objects defined in multiple feature dimensions. Instead of shifting the template away from distractors within one dimension, features in more diagnostic dimensions (in which the target is unique, or nearly unique) may then also be emphasized in the template. In the coffee mug example, this could mean that the template more strongly reflects the mug’s shape than the mug’s colour when searching on the windowsill. Two recent behavioural studies provided evidence for this idea (Boettcher et al., 2020; Lee & Geng, 2020). In these studies, participants searched for gratings defined by orientation and colour (e.g., a blue grating at 45-degree orientation). In each block of trials, one dimension (e.g., colour, but varying particular features as red or blue across trials) could be used to differentiate the target from the distractors on most trials. On a subset of trials, however, the target was not unique in the expected dimension of that particular block, but instead stood out in the other dimension (e.g., orientation). Performance was worse on these mismatching trials, suggesting that target-features within the diagnostic dimension were emphasized relative to the expected undiagnostic one.

While these two studies provide evidence that attentional templates are influenced by distractor context, they leave several important questions unaddressed. First, while coloured gratings consist of two clearly separate dimensions, distinctly represented in early visual cortex, in daily life we typically search for object shapes that have a more unified representation in intermediate and higher-level areas of the ventral visual cortex (Kourtzi & Connor, 2011). Because most target objects are not defined by a combination of two distinct low-level features, it is not clear whether attentional templates for object shapes are similarly biased by distractor context to emphasise particular shape dimensions.

Second, it is unclear *exactly how context-dependent* such adaptive biases in the template are and what mechanism underlie these changes. Virtually all previous studies required participants to search repeatedly in contexts where either the distractor features (e.g., S. I. Becker, 2010b; Geng et al., 2017; Yu & Geng, 2019) or target-distractor relations (Boettcher et al., 2020; Lee & Geng, 2020) remained stable over an experimental block, or even the whole experiment. Expectations could therefore be based on recent selection history. Previous searches dynamically shape the attentional template, without necessarily requiring that target-distractor relations are learnt in a contextualized manner. Biases based on priming, a facilitation in performance based on repeating targets or distractors over successive trials, can reflect the statistics of targets and distractors (see Kristjánsson, 2022 for a review). Notably, such priming effects have been found for repetitions of specific features (e.g. *red*; Maljkovic & Nakayama, 1994), feature relations between targets and non-targets (e.g. *spikier* S. I. Becker, 2013), but also entire feature dimensions (e.g. *overall colou*r; Found & Muller, 1996; Liesefeld et al., 2019). These biases are however typically short-lived. This is relevant as we often don’t search repeatedly for the same object in the same context, but rather search for different objects in the same context or for the same object in different contexts. In such arguably more naturalistic situations (as in our coffee mug example), target-distractor relations would have to be learnt and bias the template in a context-dependent manner to account for changing target-distractor relations across contexts. Accordingly, it should be possible to base distractor expectations on long-term memory for different contexts, rather than short-term priming mechanisms. After learning these relations, re-encountering familiar contexts may flexibly shape the attentional template even on a trial-by-trial basis.

Finally, another relevant open question is whether the findings of previous studies relied on participants having explicit knowledge of the block structure or whether distractor context effects can arise implicitly. While Lee et al. (2020) found equivalent effects of distractor context when participants were either explicitly informed about the distractor context manipulation or not, they did not assess whether participants were nonetheless aware of this in both cases. Considering that the stimulus dimensions in this experiment were clearly distinct and that the same dimension was diagnostic throughout half of the experiment, it is likely that participants were aware of the probabilities for different diagnostic dimensions and strategically used this. If templates are adapted to distractor context in daily life, this should occur relatively effortlessly and possibly implicitly.

Here, in a series of experiments, we addressed these open questions, probing when and how attentional templates are shaped by distractor context. We familiarized participants with novel, complex shapes and asked participants to search for them in different probabilistic contexts where one of the shape dimensions was diagnostic of the presence of the target. We first tested whether attentional templates for these shapes were biased by distractor expectations in a blocked context (Experiments 1 and 2) and assessing awareness of the manipulation (Experiment 2). Then, we investigated whether distractor expectations would also influence templates when context varied on a trial-by-trial basis (Experiments 3 and 4). To summarise the results, across experiments, we found flexible attentional templates for complex shape stimuli, shaped by distractor expectations in blocked contexts, independently of awareness. Further, we found evidence for context-dependent attentional templates, dynamically adjusted to context expectations on a trial-by-trial basis (Experiment 4).

### Transparency and Openness

Experimental stimuli, code and data are available on the porject’s OSF page (https://osf.io/tgu8r). All analyses were done using Matlab (2019a), the measures of effect size toolbox (Henschke, 2002) and JASP (JASP Team, 2022).

Design and analyses of Experiments 2B, 3 and 4 were preregistered on AsPredicted: https://aspredicted.org/bg5qa.pdf (Exp 2B), https://aspredicted.org/e9zs9.pdf (Exp 3), https://aspredicted.org/y4tq4.pdf (Exp 4).

To determine sample size, a priori power analyses were run using G*Power (Faul et al., 2007).

## Experiment 1 – Diagnostic Target Dimensions are Emphasized in Blocked Distractor Contexts

In the first experiment, we tested the effect of blocked distractor context in which one target dimension was more discriminative than another when searching for complex 2D shapes. We created shapes varying along two dimensions: rectilinearity and orientation, jointly contributing to the outer form of the shape (see Figure 1)^1^. In addition, to increase similarity to real-world objects, all shapes were given names and additional elements that made them unique, characterizing them as individuals. While these stimuli were still controlled and simpler than real-world objects, they were designed to evoke an integrated representation. After familiarizing themselves with the target shapes and learning their names, participants searched for these shapes in displays in which either the target’s rectilinearity or orientation was unique. To create predictable distractor contexts, unique rectilinearity and orientation trials were grouped in rectilinearity or orientation context blocks (with 80% of trials within a block belonging to this trial type). Participants were not explicitly informed about this. Due to the blocked structure, we could compare performance for the same trial type when it either matched or mismatched distractor expectations based on the block they were presented in. Importantly, displays on context-matching and -mismatching trials were visually identical, the only difference being the block context in which they appeared.

**Figure 1.**
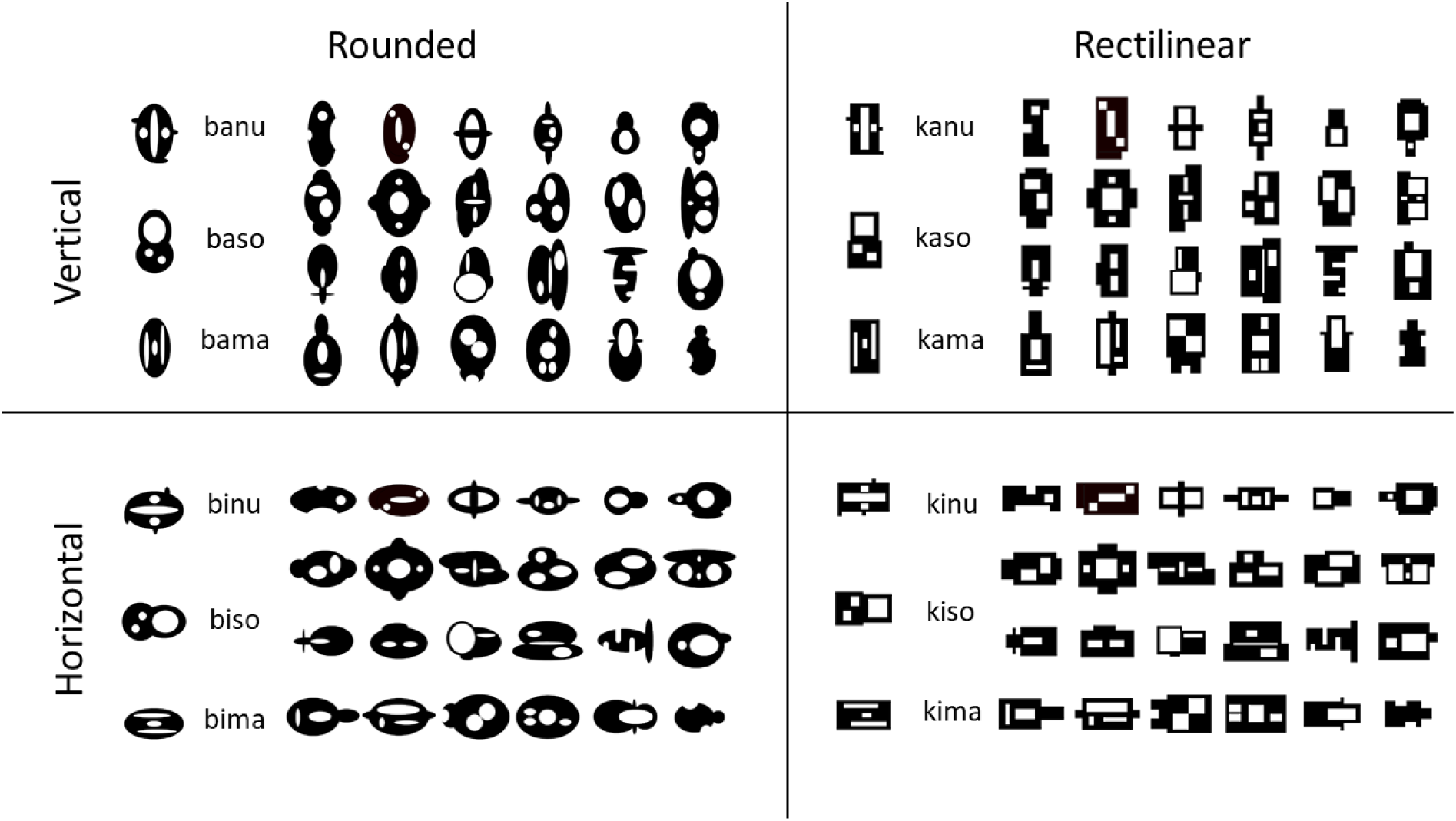
Overview of all Target and Distractor Shapes Note. Target shapes are the named shapes in the leftmost column of each quadrant, all other shapes are distractors.

We chose to cue the shapes by name instead of presenting them visually prior to search. This required participants to rely on an internally generated memory representation based on the learned associations between shape names and visual features to create a template. Arguably, generating a template in this top-down manner more closely resembles real-life search than visually cued search. However, compared to visually cued targets (e.g., Lee & Geng, 2020), this may lead to less flexible templates (see Boettcher et al., 2020 for a discussion). This is because, first, there is no external visual information of which different features or feature dimensions could be selectively encoded or maintained in visual working memory (e.g., Niklaus et al., 2017; Park et al., 2017); and, second, the memory representation on which the template is based has to be repeatedly retrieved across different contexts and may therefore not be biased by a particular context itself. Nevertheless, there is evidence that such associative templates can still be biased by context (Boettcher et al., 2020).

### Methods

#### Participants

A total of 43 participants were recruited via Prolific (https://www.prolific.co) and took part in the online experiment. Thirty-four participants (mean age: 28.08, sd: 4.91; 22 female) were included in the analysis. 7 participants were excluded due to low task performance (< 55% accuracy, with chance level being 50%) and two further participants were excluded because they relied on a strategy based on shape familiarity without using the shape name cue in the search task: on trials where the cued shape was presented together with another distractor taken from the set of 12 target shapes on the other side of the display, their selection of the cued shape was not above chance (as determined by a binomial test with α = .05). We excluded these participants because there was no evidence that the participants created a feature-specific template based on the cue (and which may then be modulated by distractor context expectations). Instead, they appeared to respond indiscriminately to any of the 12 familiar target shapes.

The final sample size of 34 was determined a priori. Assuming an effect size of about *η_p_²* = .20 for the critical main effect of context match (a lower estimate consistent with results from Lee & Geng (2020)), this allowed for 80% power (with α = .05) .

All participants provided informed consent and were paid £4.50. The study was approved by the Radboud University Faculty of Social Sciences Ethics Committee (ECSW2017-2306-517). All data were collected in 2021.

#### Procedure

The experiment was divided into a short shape-name training phase of at least 3 blocks (8 trials each) and 6 blocks (40 trials each) of the search task, taking around 35 minutes in total.

During the training phase participants learnt the names of 12 target shapes (see Figure 1), such that these names could later be used as cues in the search task. These individual shapes could be subdivided into 4 subgroups, sharing the same orientation (vertical or horizontal) and rectilinearity (rectilinear or rounded form). To support learning, all shape names provided information about the appearance of the shape, with the first letter (k or b) indicating its rectilinearity, the second letter (i or a) its orientation and the last two letters (ma, nu or so) the individual shape within that group. For each training block, participants first saw a display showing the 4 shapes and shape names belonging to the same individual (e.g., the baso, kaso, biso and kiso shapes). Then they completed 8 self-paced trials in which one of the names was presented and the correct shape had to be selected out of the four individuals in the display. If 6 out of 8 trials were answered correctly the next four individuals were presented, otherwise the training block was repeated.

After successfully completing training, participants proceeded to the search task (see Figure 2). At the start of each trial, one of the 12 target shapes was cued by name. After a delay period, a search display consisting of 6 shapes appeared for 220 ms. Participants had to report on which side of the display the target had appeared by using the left and right arrow keys of their keyboard, before receiving feedback via a coloured dot (either red or green) appearing at fixation. From the perspective of the participant, the task was always to find the cued shape. What was not mentioned to them was that on every trial, the target was either unique in its rectilinearity (e.g., a vertical rectilinear shape among rounded vertical and horizontal shapes) or orientation (e.g., a vertical rectilinear shape among horizontal rectilinear and rounded shapes) from all other distractors. In total, there were 120 unique orientation and 120 unique rectilinearity trials. Trial type was blocked with 80% validity, yielding rectilinearity and orientation context blocks (with 32 trials within a block of one trial type and 8 of the other). Mismatching context trials in the rectilinearity context block therefore had a unique orientation and vice versa. Importantly, within a block, there was not a specific feature (e.g., a horizontal shape) that was more common for either target or distractors, as all vertical and horizontal, rectilinear and rounded shapes were overall equally frequent. The difference between blocks was which targets and distractors were paired together in specific displays, making a dimension (e.g., orientation) but not any specific feature (e.g., horizontal) diagnostic. This design avoided low-level adaptation and intertrial priming of specific features (Maljkovic & Nakayama, 1994).

**Figure 2.**
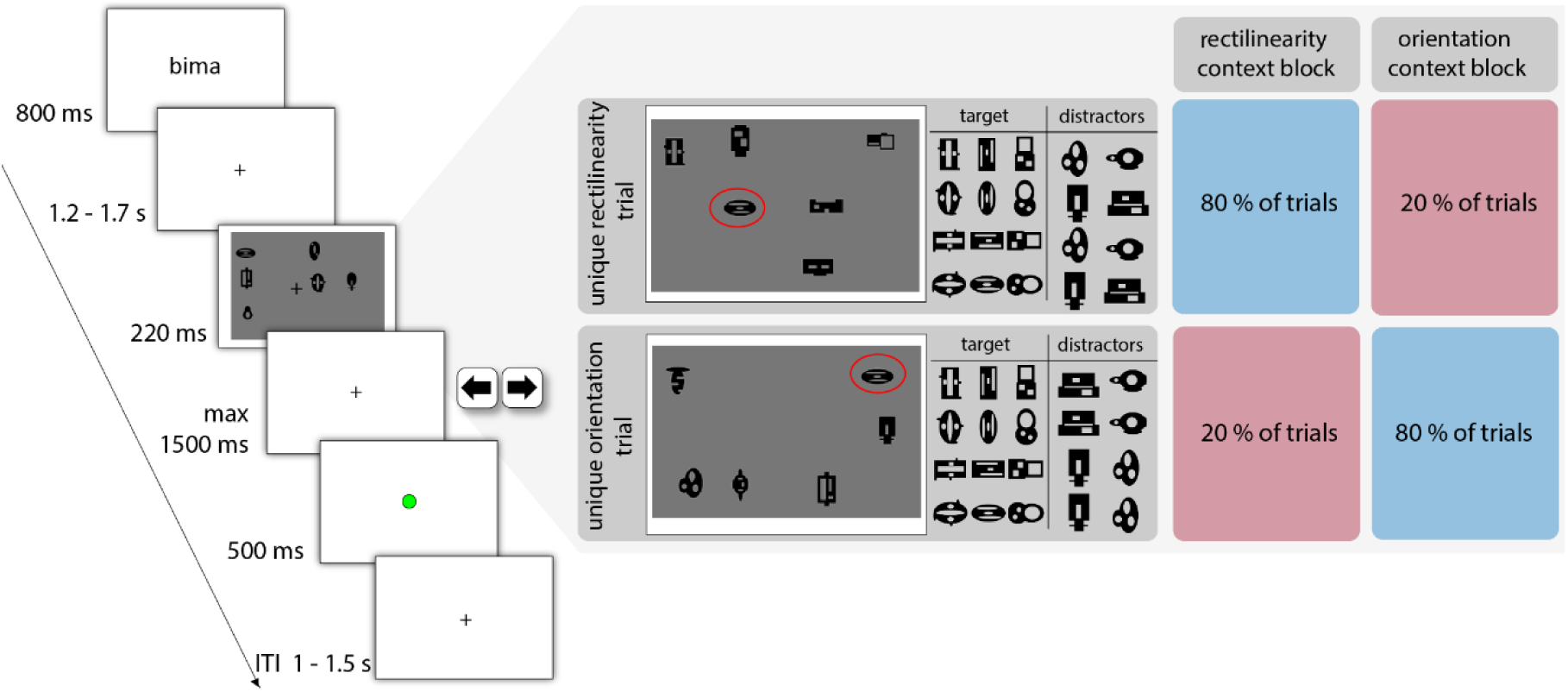
Timeline of a Trial and Design Overview for Experiment 1 Note. At the start of each trial, participants saw a shape name cue, then a search display containing the target and other shapes appeared briefly and they indicated on which side the target had appeared. On half of the trials the target had a unique rectilinearity and on the other half a unique orientation. This discriminative dimension was blocked, with 80% validity. The same search displays were thus presented in blocks in which their diagnostic dimensions matched (red) or mismatched (blue) the current block context. The target is highlighted in the example displays for illustration in this figure only. Overall, all distractors were equally frequent in both types of trials, but paired with different targets (only some example distractors are shown here).

Target orientation and rectilinearity were counterbalanced within each block, while individual target shapes and the side on which the target appeared were counterbalanced across all 240 trials of the experiment. Block type (rectilinearity or orientation context blocks) alternated and the first 8 trials of a block were always valid (i.e., belonging to the dominant trial type of that block). The initial block context (rectilinearity or orientation) was randomized.

#### Stimuli & Setup

A total of 108 novel, black shapes were created using Adobe Illustrator (see Figure 1). These shapes could be grouped into 4 categories based on their rectilinearity (rectilinear or rounded base shape) and orientation (upright or horizontal), comprising 27 individuals each. All individual shapes had a different configuration of local elements (e.g., small holes/protrusions of varying sizes and positions) and also differed slightly in their size and width-height ratio. The arrangement of these configurations was kept across shape groups, meaning that highly similar individual shapes appeared in all 4 shape groups. Shapes were created by adding these local elements to an oval or rectangular base shape, exchanging all rectangles by circles or ovals and/or rotating the resulting shapes to create similar shape exemplars across groups. Three shapes of each group were chosen as target shapes.

To create search displays, 5 distractor shapes and the target shape were placed on a grey background aligned on an invisible 3 x 4 grid, with the constraint that 3 shapes should appear on each side of the fixation cross (Figure 2). Random jitter (+/- 0.2 x the shape size) was added to the grid positions. In each display, distractors were chosen such that the target was always unique in one of its dimensions but shared features with some distractors in the other dimension, to reduce pop-out of the target. Of the 5 distractors, 3 had the same feature value (e.g., horizontal orientation) of the target in the non-discriminative dimension. Two distractors did not share any feature with the target and were presented on the same display side as the target. This allowed to distinguish responses to the target and to the most similar distractors. For a given target shape, distractors in the display could also include other target shapes from other shape groups, but these were not more likely than any other non-target distractor.

The experiment was programmed in Psychopy (Peirce et al., 2019) and hosted on Pavlovia (https://pavlovia.org). All participants took part in the experiment using their own computer. At the beginning of the experiment, their screen resolution was determined by asking them to rescale a credit card picture on the screen to the physical size of their credit card and they were asked to sit at about an arm’s length (∼57 cm) from their screen (Li et al., 2020). Assuming this viewing distance, the search displays subtended around 20 x 15 dva.

#### Data Analysis

For our analyses, accuracy and reaction were combined into a single measure, the linear integrated speed accuracy score (LISAS; Vandierendonck, 2017). This provided us with a single performance measure that is sensitive to effects in accuracy and RT in the same direction, accounting for additional variance in performance compared to its two individual components. Individual results for accuracy and RT for every experiment are included in the Appendix. Additional figures for accuracy and RT results can also be found in the supplementary materials (figures S1 – S4).

The LISAS is a linear combination of RT and error rates, the latter scaled by the standard deviation of both measures, thus increasing reaction times as a function of error rates. It is defined as: 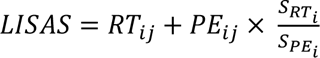, with 𝑅𝑇_𝑖𝑗_ being the mean (correct) response time of participant 𝑖 in condition 𝑗, 𝑃𝐸_𝑖𝑗_ participant 𝑖’s error rate in condition 𝑗, 𝑆_𝑅𝑇𝑖_ the standard deviation of response time (for participant 𝑖) and 𝑆_𝑃𝐸𝑖_ the standard deviation of responses (coded as 0’s and 1’s for correct/incorrect) of this participant.

To test for effects of distractor context expectations, we analysed performance on the unique rectilinearity and orientation trials as a function of the block context using a 2 (trial type: unique rectilinearity, unique orientation trial) x 2 (context match: presented in matching, mismatching context block) repeated-measures ANOVA.

Context matching effects for individual dimensions were followed up with t-tests (2-sided, α = .05). Bayes factors for simple t-contrasts and the main effects/interactions were calculated as Bayesian one-sample t-tests for the respective difference scores between conditions or condition averages using the JASP default settings (Cauchy prior with scale 0.707).

### Results

Trials with reaction times below 150 ms and those +/- 3 standard deviations away from participant’s mean RT on correct trials were excluded, resulting in rejection of 1.26% (sd 0.93) of trials.

Overall accuracy was 82.25% (sd 8.52), overall RT was 696.83 ms (sd 82.85).

As hypothesized, participants performed better when distractor context was expected, i.e. when a unique rectilinearity/orientation trial appeared in the matching compared to mismatching block context (Figure 3; main effect of context match: *F*(1,33) = 7.84*, p* = .009, *η_p_²* = .19, *BF_10_* = 4.98).

**Figure 3.**
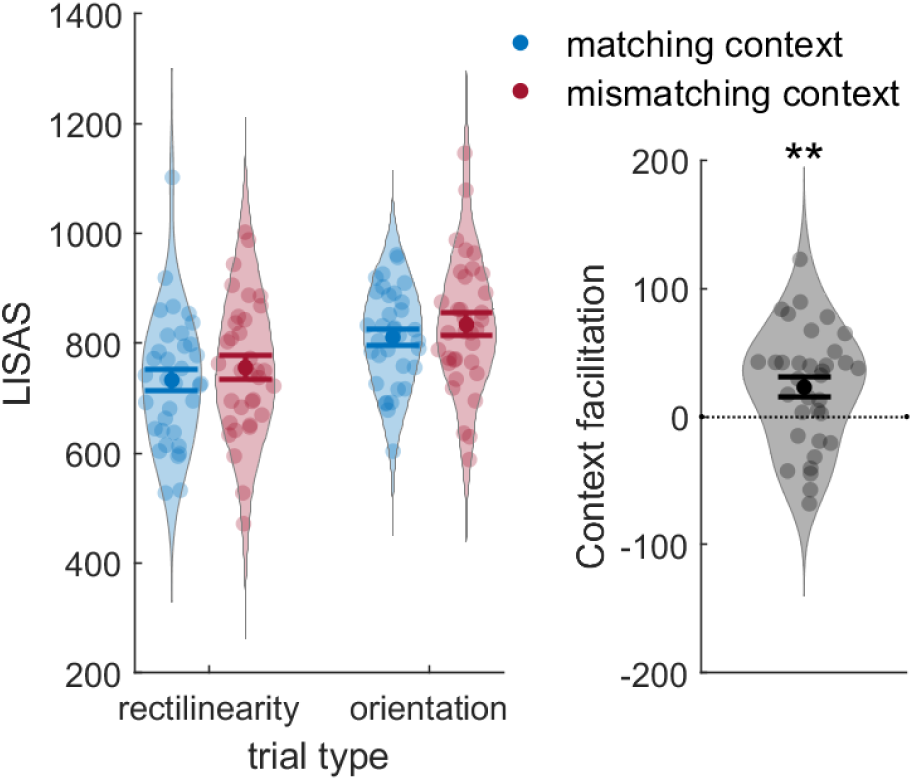
Results of Experiment 1 Note. Performance on both trial types as a function of block context (left) and context facilitation (LISAS mismatching – matching context) averaged across trial types (right). All error bars are SEM.

In addition, participants were overall better on the unique rectilinearity trials (main effect of trial type: *F*(1,33) = 50.92 *p* < .001, *η_p_²* = .61, BF_10_ > 3 x 10^5^). This suggests that the target was generally easier to find when it had a unique rectilinearity than when it had a unique orientation. However, context expectations influenced search equally for both trial types (no significant interaction between trial type and context match: *F*(1,33) = 0.00, *p* = .99, *η_p_²* < .001, BF_10_ = 0.18; context match rectilinearity unique trials: t(33) = 1.51, p = .14, d_z_ = 0.26, BF_10_ = 0.51; orientation unique trials: t(33) = 1.43, p = .16, d_z_ = 0.24, BF_10_ = 0.46).

### Discussion

Experiment 1 showed that search performance was influenced by distractor context expectations. Searching for the same target shapes in different block contexts, participants performed better when the diagnostic target dimension matched their expectations compared to when it did not, even though the search displays were identical in these conditions. These findings provide an important extension of previous work (Lee et al., 2019; Boettcher et al., 2020), showing that probabilistic distractor context influences search performance for complex shapes with individual features and names. Notably, these biases also arose relatively quickly, as each block only had 32 valid trials and learning of distractor context could only be based on briefly flashed displays with heterogeneous distractors differing in the non-diagnostic dimension.

We interpret these results as reflecting a change in the attentional template, with a different template being used for the same object in different contexts based on distractor expectations. As we used name cues, this bias cannot be explained in terms of selective encoding or maintenance (Niklaus et al., 2017; Park et al., 2017) of external visual information into working memory. Thus, we show that internally-generated templates for complex shapes, as used in naturalistic search, are flexibly shaped by distractor expectations.

## Experiment 2 – Probing Biases in the Target Template

Our interpretation of the results of Experiment 1 is that the attentional template for the shapes was modulated by distractor expectations, and that the target feature in the relevant dimension was thereby emphasized over the irrelevant dimension (Lee & Geng, 2020). However, another possibility might be that this representation of the target shape itself did not change, but rather that participants were learning about some property of the entire display, without changing their template for the target shape as such.

For example, despite attempts to make the displays homogeneous and the target not pop out, our target shape was by definition still a singleton in one dimension. Participants could therefore have learnt to look for orientation or rectilinearity singletons depending on the block and not created/modulated shape-specific target templates. This is especially relevant given that singleton search is facilitated when the target-defining dimension repeats across trials (e.g. Found & Müller, 1996). Indeed, according to the dimension weighting account of attention (see Liesefeld et al., 2019 for a review), feature dimensions can be weighted differently when information is integrated across dimensions into a single spatial priority map (see S. I. Becker, 2010a; S. I. Becker et al., 2014 for a post-selectional account for dimension-weighting effects in these studies). These effects are however different from a change in the attentional template to emphasize a specific feature, as singleton search only requires to find the odd-one out without requiring a specific template. Because this account is based on saliency or contrast, dimension weighting also applies equally to all features in one dimension (see also Lee & Geng, 2020).

Along similar lines, based on which distractors appeared together in displays of different contexts, participants could also have learnt to efficiently group and suppress distractors with a common rectilinearity in the rectilinearity context blocks and a common orientation in the orientation block.

To test whether our predictive distractor context indeed changed the shape-specific attentional template, we adapted our paradigm to include 2-shape probe-trials, in which the target was presented along with the same individual of another shape group (shapes sharing similar local elements, but always differing in one dimension, e.g., a kaso and a kiso shape). These trials were presented intermixed with the regular 6-shape search displays, for which the discriminative dimension was always consistent within a block. If the target template was indeed shaped by context, selecting the target should be easier based on the discriminative dimension of the 6-shape displays in this block. We would therefore expect to replicate the results of Experiment 1 with the 2-shape displays. If, instead, participants learnt to select a specific singleton or to suppress certain distractor groups in the display, this learning should not transfer to the 2-shape trials and performance on these trials should therefore be independent of context.

In addition, we were also interested in whether our results reflected explicit knowledge of distractor context. Previous studies provided evidence that different aspects of distractor context, including co-occurrence between distractors (Thorat et al., 2022), their spatial arrangements in contextual cueing paradigms (Chun & Jiang, 1999; Spaak & de Lange, 2020), and the shape of their underlying feature distribution (Hansmann-Roth et al., 2021) can be learnt implicitly, without awareness of these regularities. Here, we introduced two questions with ratings at the end of the experiment, asking participants whether they had become aware of the distractor context manipulation and whether they had used it strategically. We then related these ratings to context facilitation.

### Methods

#### Participants

Data from 34 participants (18 female; mean age: 28.44, sd: 4.93) were included for the original experiment. 2 participants were replaced due to relying on a familiar shape strategy rather than using the shape name cue, as described in Experiment 1. As for Experiment 1, sample size was determined a priori and allowed to detect a main effect of context match with effect size of *η_p_²* = .20 with 80% power (α = .05).

After the original experiment (Experiment 2A), we ran a higher-powered, exact replication (Experiment 2B). 80 new participants were included (45 females, 1 person who did not indicate their gender; mean age: 29.18, sd: 4.12). Five participants were replaced due to overall low accuracy (<55%) and an additional one because they did not use the shape name cue. Based on this sample size, we were able to detect a smaller effect (about half of the effect size in Experiment 1) with 80% power.

All participants were recruited via Prolific (https://www.prolific.co), provided informed consent prior to participation and received £5.00. The study was approved by the Radboud University Faculty of Social Sciences Ethics Committee (ECSW2017-2306-517). All data were collected in 2022.

#### Procedure

Participants were first familiarized with the shapes and their names using the same training procedure as in Experiment 1. Afterwards, they completed 10 self-paced practice trials and 6 blocks (64 trials each) of the actual search task.

On each trial, participants saw a shape name cue, followed by a search display containing either 6 or 2 shapes. Regardless of the number of shapes shown, participants always indicated on which side of the display (left or right) the target shape had appeared. Six-shape displays appeared on 3 out of 5 trials (40 trials per block) and within a block, all had the same discriminative dimension, creating the block context in which the 2-shape displays were presented (Figure 4). The 2-shape displays contained the target shape as well as the same individual shape from another target group (i.e., differing from the target shape in either rectilinearity or orientation, but sharing the other dimension and having similar other local elements) on the other side of the fixation cross.

**Figure 4.**
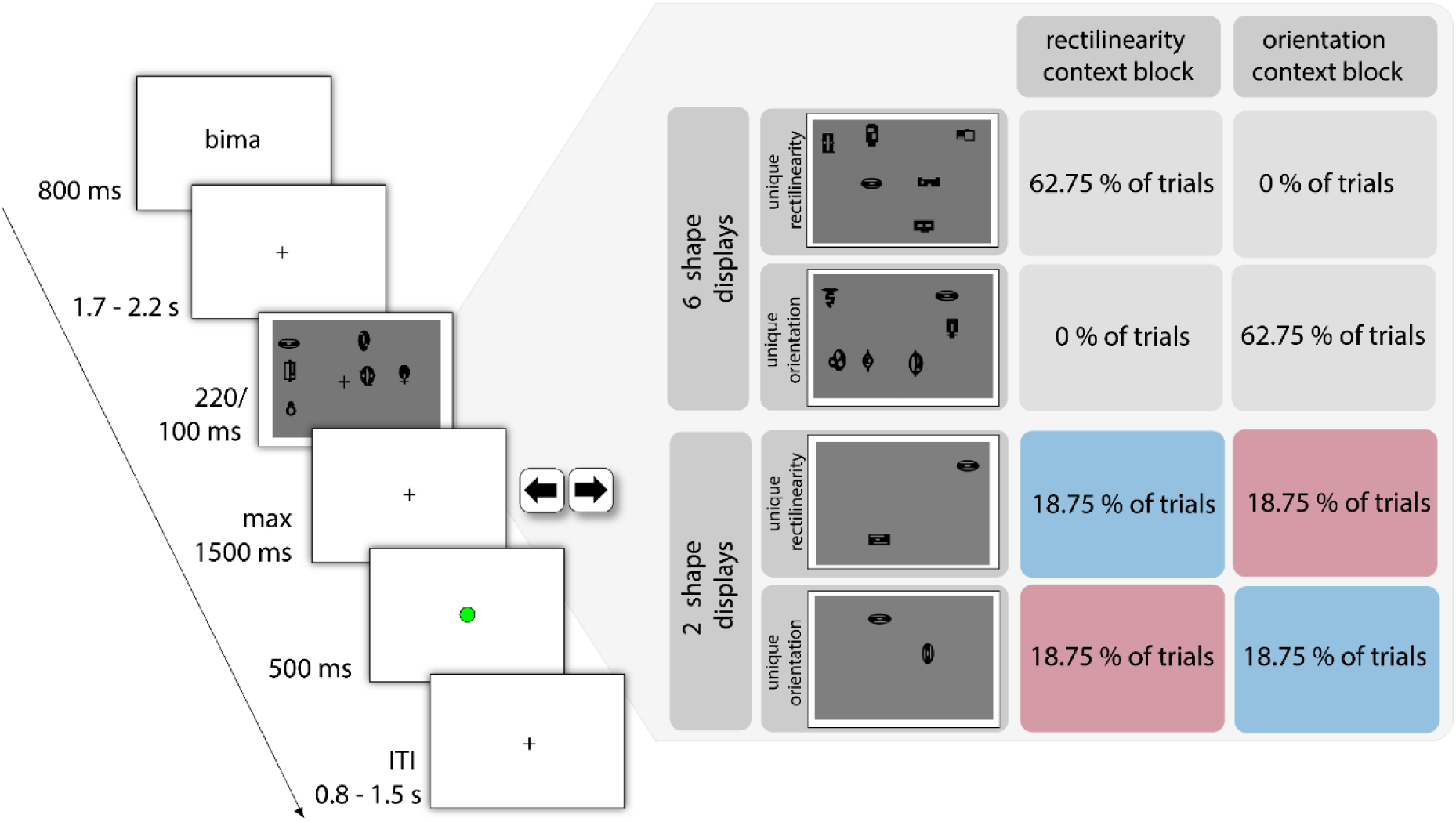
Timeline of a Trial and Design of Experiment 2A and 2B Note. Shape targets were cued by name and had to be searched for in displays of 2 or 6 shapes. 6-shape displays within one block all had the same diagnostic dimension and created the context in which 2-shape displays were presented. Those always contained the target and the same individual taken from another shape group, differing either in rectilinearity or orientation and therefore matching (blue) or mismatching (red) the current context.

Independent of the block context, on half of these 2-shape trials, the target shape had a unique rectilinearity; on the other half, it had a unique orientation. Collapsed over 6- and 2-shape trials, the diagnostic dimension was 81.25% valid within a block, similar to Experiment 1.

In an attempt to equalize difficulty across display types, 2-shape displays were presented for 100 ms only, compared to 220 ms for the 6-shape displays. Following their response, participants received feedback via a red/green circle presented at fixation.

After completing the experiment, participants were told about the distractor context manipulation and asked to report 1) whether they had noticed that all 6-shape trials in a block had a unique discriminative dimension, and 2) whether they had actively used this, by providing ratings on 5-point scales.

Target orientation and rectilinearity was counterbalanced within blocks, while individual target shapes and display side of the target were counterbalanced across the whole experiment. All factors were counterbalanced separately for each display type. Block context alternated and the block context participants started with was randomized.

#### Stimuli & Setup

The 6-shape displays were created as in Experiment 1. For the 2-shape displays, the target shape was placed on one side of the display, and the same-individual shape from another shape group (sharing either rectilinearity or orientation and other local elements with the target) on the other side. As for the 6-shape displays, an invisible 3 x 4 grid was used to place the shapes.

All participants took part in the experiment on their own device. The experiment was coded using Psychopy and hosted on Pavlovia (https://pavlovia.org).

#### Data Analysis

As for Experiment 1, we conducted a 2 (trial type: unique rectilinearity, unique orientation trial) x 2 (context match: presented in matching, mismatching context block) repeated measures ANOVA on the LISAS of the 2-shape trials.

In addition, we compared the effects of context match on the 2-shape trials of Experiment 2B with the context match effects of Experiment 1 using a 2 (trial type: unique rectilinearity, unique orientation trial) x 2 (context match: presented in matching, mismatching context block) x 2 (Experiment: Experiment 1, 2B) repeated measures ANOVA, with Experiment as between participant factor.

Context matching effects for individual dimensions were followed up with t-tests (2-sided, α = .05). Bayes factors for simple t-contrasts and the main effects/interactions were calculated as Bayesian one-sample t-tests for the respective difference scores between conditions or condition averages using the JASP default settings (Cauchy prior with scale 0.707). For comparisons between experiments or groups, Bayesian independent sample t-tests were computed instead.

### Results

#### Experiment 2A

We excluded trials in which response times were below 150 ms or +/- 3 standard deviations away from the participant’s mean correct RT for the respective display type, on average leading to a rejection of 1.56% (sd 0.87) of trials across display types.

Accuracy for the 6-shape trials was 89.35% (sd 6.97) and mean correct RT was 726.37 ms (sd 84.44). For the 2-shape trials, accuracy was 90.52% (sd 8.61) and mean correct RT was 670.18 ms (sd 84.48).

All subsequent analyses focused on the 2-shape trials, as the 6-shape trials did not include context mismatch trials.

Overall, there was a slight improvement in performance when the target’s diagnostic dimension was expected based on block context, but this was not significant (Figure 5; main effect of context match: *F*(1,33) = 1.78, *p* = .19, *η_p_²* = .05, *BF_10_* = 0.42). As previously, trials in which the target’s rectilinearity was diagnostic were generally easier (trial type: *F*(1,33) = 34.77, *p* < .001, *η_p_²* = .51, *BF_10_* >1000). In this experiment, the effect of context expectations also differed between trial types (context match x trial type: *F*(1,33) = 7.39, *p* = .01, *η_p_²* = .18, *BF_10_* = 4.20): for rectilinearity unique trials performance improved when they were presented in a matching context (*t*(33) = 2.64, *p* = .01, *d_z_* = 0.45, BF_10_ = 3.54), but this was not the case for unique orientation trials (*t*(33) = -1.43, *p* = .16, *d_z_* = -0.25, BF_10_ = 0.46).

**Figure 5.**
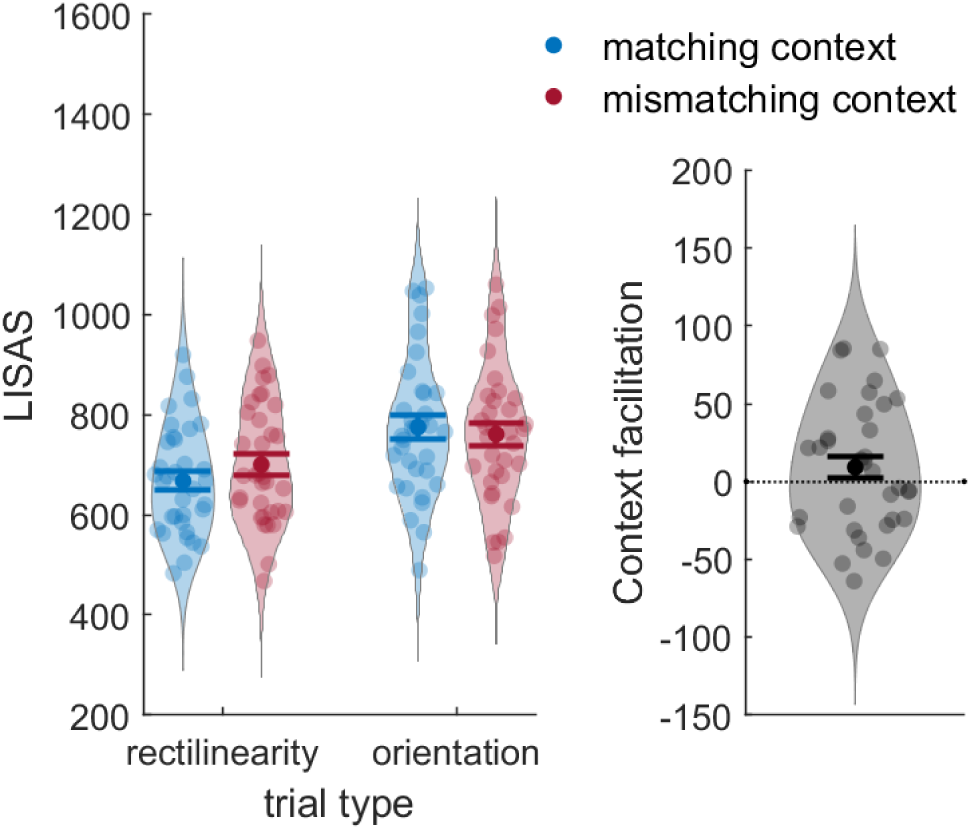
Results of Experiment 2A Note. Performance on the 2-shape trials for both trial types as a function of block context (left panel) and context facilitation (LISAS mismatching – matching context) averaged across trial types (right). All error bars are SEM.

We then tested for overall awareness of our context manipulation and its relation to context facilitation. Twenty-eight participants provided an awareness rating, ranging from 1 (I didn’t notice this at all) to 5 (it was very obvious). The mean value was 3.00 (sd 1.39), with 6 participants indicating they had not noticed the relevant dimension being consistent within blocks at all (corresponding to a rating of 1) and 12 participants giving it a rating >= 3.

For the second rating, regarding whether participants actively made use of context knowledge, we excluded participants who had responded with 1 for the awareness rating (as they could not provide meaningful answers to this question), resulting in 22 ratings. The rating ranged from 1 (I didn’t use this at all) to 5 (I used this a lot and prepared differently depending on the blocks). The mean rating was 2.95 (sd 1.39), with 3 participants indicating they had not made use of it at all (rating of 1) and 12 providing a rating >= 3.

We correlated both of these ratings with participant’s context facilitation (LISAS mismatching – matching context, averaged across both trial types). Context facilitation did not correlate with either the awareness (*ρ* = - .08, *p* = .69) or the usage rating (*ρ* = -.13, *p* = .56). This was also true when correlating only context facilitation in unique rectilinearity trials with awareness (*ρ* = .02, *p* = .91) or usage ratings (*ρ* = -.13, *p* = .56).

#### Experiment 2B

We excluded anticipatory and delayed responses for every participant, as preregistered and described for the original experiment, resulting in an average loss of 1.54% (sd 0.77) of trials.

For the 6-shape trials, accuracy was 89.16% (sd 7.47) and RT was 731.06 ms (sd 101.02). For the 2- shape trials, accuracy was 90.79% (sd 8.28) and RT was 666.48 ms (sd 96.67).

The analysis was the same as Experiment 2A. In line with our hypothesis, we found that performance improved when the diagnostic dimension matched participants’ expectations (Figure 6; main effect of context match (*F*(1, 79) = 9.98, *p* = .002, *η_p_²* = .112, *BF_10_* = 11.70). Our higher-powered replication thus confirmed that the target template was indeed modulated by distractor expectations.

**Figure 6.**
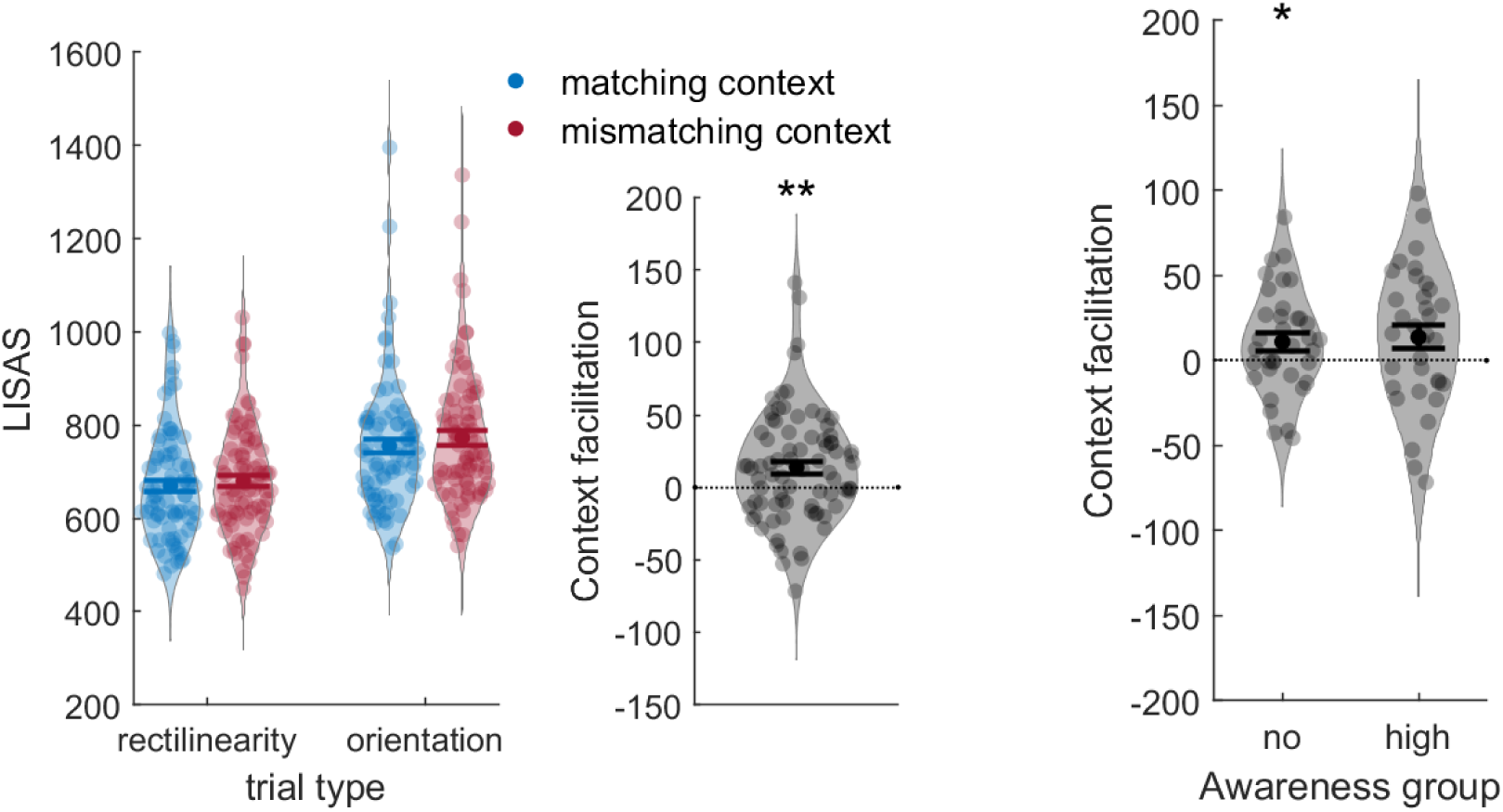
Results of Experiment 2B Note. Performance on the 2-shape trials for both trial types as a function of block context (left panel) and context facilitation (LISAS mismatching – matching context) averaged across trial types (middle panel). The right panel shows context facilitation for Exp 2A & 2B combined, split by participants’ reported awareness of the context manipulation. All error bars are SEM.

Again, performance was overall higher when rectilinearity was the diagnostic target dimension (trial type: *F*(1,33) = 95.76, *p* < .001, *η_p_²* = .55, *BF_10_* > 2x 10 ^12^). Unlike Experiment 2A, the effect of context expectations was independent of which dimension was diagnostic (context match x trial type: *F*(1,33)= 0.28, *p* = .60, *η_p_²* = .00, *BF_10_* = 0.14; context match unique rectilinearity trials: *t*(79) = 1.32, *p* = .19, *d_z_* = 0.15, *BF_10_* = 0.28, unique orientation trials: *t*(79) = 2.13, *p* = .04, *d_z_* = 0.24, *BF_10_* = 1.04).

We also tested whether context facilitation on the 2-shape trials differed from the one found for 6-shape trials in Experiment 1 in an exploratory analysis. Even though these differed in some aspects (most importantly that the 2-shape trials were generally easier, but also the number of trials), context facilitation did not differ across experiments (main effect of experiment: *F*(1,112) = 7.68, p = 0.007, *η_p_²* = .06, *BF_10_* = 6.08; no interaction between context match x Experiment: *F*(1, 112) = 1.10, *p* = .30, *η_p_²* = .01, *BF_10_* = 0.26).

In addition to the main preregistered analysis, awareness and usage ratings were also collected for this experiment. We collected 76 awareness ratings, having a mean value of 2.34 (sd 1.34), with 29 subjects indicating that they had not noticed this at all (giving a rating of 1) and an equal number of subjects giving it a rating >= 3. For the usage rating, 47 ratings were included and the mean value was 2.89 (sd 1.27), with 28 ratings >= 3.

As in Experiment 2A, neither the awareness, nor the usage rating correlated with the context facilitation across participants (awareness rating: *ρ* = .08, *p* = .48, usage rating: *ρ* = - .09, *p* = .55), suggesting that context facilitation was not related to explicit knowledge about the distractor context.

To test this more explicitly, we pooled data from Experiment 2A and 2B to repeat our analysis of distractor context effects separately for participants who reported not having noticed the distractor manipulation (awareness rating = 1) and those who found it relatively obvious (awareness rating >= 4) in an exploratory analysis. Most importantly, we found that in the no-awareness group, performance still improved when the diagnostic dimension was expected based on context

(main effect of context match; no awareness group: *F*(1,33) = 4.53, *p* = .04, *η_p_²* = .12, *BF_10_* = 1.33; high awareness group: *F*(1,34) = 4.00, *p* = .05, *η_p_²* = .11, *BF_10_* = 1.07), and the effect of context expectations was not stronger for the high awareness group (*t*(67) = - 0.31, *p* = .75, *d_z_* = 0.05, *BF_10_* = 0.26). Also the correlations of context facilitation with awareness and usage ratings were not significant across this larger sample (awareness rating: *ρ* = .01, *p* = .96, usage rating: *ρ* = - .13, *p* = .29).

### Discussion

Experiment 2 provided additional evidence that distractor expectations shapes the representation of target features: selecting the target in the 2-shape trials based on its rectilinearity was easier in the rectilinearity blocks than orientation context blocks and vice versa. By intermixing 2-shape trials in the blocks we were able to show that distractor context affected the attentional template, and therefore the representation of the target shape, rather than more efficient grouping of distractors or singleton detection.

In Experiment 2A (but not Experiment 2B), we observed a difference in context facilitation across diagnostic dimensions, with the modulation by distractor expectations being stronger for the rectilinearity than the orientation dimension. A similar kind of asymmetry was found in previous studies (Boettcher et al., 2020; Lee & Geng, 2020), where colour was more consistently modulated by context compared to orientation. One interpretation is that feature dimensions which generally allow for easier discrimination (and the target can be more easily selected on their basis, as rectilinearity in our task, or colour in others) are more accessible and likely to be further emphasized or deemphasized based on context.. In addition, if distractors are sharing target features in this dimension, they may be particularly distracting and require adjusting the template, more so than for overall less prominent dimensions. For example, if target rectilinearity is more strongly represented than orientation in the template by default, especially the distractors sharing target rectilinearity in the unique orientation trials would hamper performance (in line with the overall worse performance observed on these trials). Encountering many such distractors in the unique orientation blocks would require adjusting this default, and decreasing the reliance on rectilinearity. Therefore performance on the rare unique rectilinearity trials in orientation blocks may be worse than in the rectilinearity unique blocks, leading to a context facilitation effect for rectilinearity trials.

Was this change in the template due to deliberate and explicit processes or could it occur more implicitly? The awareness ratings suggest that our manipulation became noticeable to several participants at some point during the experiment, and a subset of those also reported actively using knowledge of the likely discriminative dimension. However, both ratings did not correlate with the magnitude of context facilitation across participants. Further, we also found context facilitation for those participants who reported not having noticed any distractor regularities, which was equal in magnitude to the facilitation shown by participants who did notice them. While this small effect should be interpreted with care and further replicated, modulation of the attentional template by distractor expectations may therefore not depend on participants noticing and deliberately using distractor regularities, similar to the implicit learning of other types of distractor regularities (Chun & Jiang, 1999; Hansmann-Roth et al., 2021; Spaak & de Lange, 2020; Thorat et al., 2022).

## Experiment 3 – Adjusting to Distractor Expectations on a Trial-by-Trial Basis

Having established that expected discriminative target features could be emphasized in the attentional template based on a blocked context, in Experiment 3 we tested whether the attentional template is also updated when distractor contexts are familiar but changing frequently, as would be necessary in many daily-life situations (e.g., searching for your keys in different rooms or on different tables). This would be evidence for a flexible top-down bias based on long term memory for respective contexts, independent of priming mechanisms.

In this way we could test whether target-distractor relations were learnt separately for different contexts and thus whether distractor expectations modulated search performance in a truly context-dependent manner. Previous research has shown that some statistics of targets or distractors, e.g. recent colour statistics within object categories (Kershner & Hollingworth, 2022) or features associated with rewards (Anderson, 2015) are learnt and used in that way, with context cues modulating the attentional template on a trial-by-trial basis. Furthermore, there is also neural evidence that context cues can be used to tune sensory gain of orientation-selective neurons in visual cortex away from distractor orientations on a trial-by-trial basis (Scolari et al., 2012).

To provide context cues, we paired the two different contexts (either unique rectilinearity or orientations as frequent diagnostic dimension) with two distinct background colours. To facilitate learning of the colour-context associations, the distractor context was blocked for the first half of the experiment (as in Experiments 1 and 2). Afterwards, distractor context changed randomly on every trial.

Our design now required participants to not only learn regularities in diagnostic dimensions, but also to associate these with different colour cues, which is more challenging. Since the main focus of this study was not on whether these additional associations could be learnt implicitly but rather on whether they were being used after learning, we now informed participants about the meaning of those colour cues.

### Methods

#### Participants

80 participants (52 female, 1 non-binary person; mean age: 28.63, sd: 4.17) were included in this experiment. 5 additional participants were replaced due to low accuracy (<55%). We assumed the effect of context expectations may be reduced in the interleaved condition, and this sample size allowed us to detect main effect of context match effect about half as large as the blocked condition in Experiment 1 with 80% power (*η_p_²* > .095, with α = .05).

All participants were recruited via Prolific (https://www.prolific.co), provided informed consent and received £5.00 for their participation. The study was approved by the Radboud University Faculty of Social Sciences Ethics Committee (ECSW2017-2306-517). All data were collected in 2022.

#### Procedure

The overall procedure was similar to Experiment 1 (Figure 7). Participants first completed the same shape-name training before starting the search task. The search task was explained to them, including the distractor context manipulation and that the background colour of the search display provided information about the likely upcoming distractor context. Before the start of the search task, participants also completed 10 self-paced practice trials.

**Figure 7.**
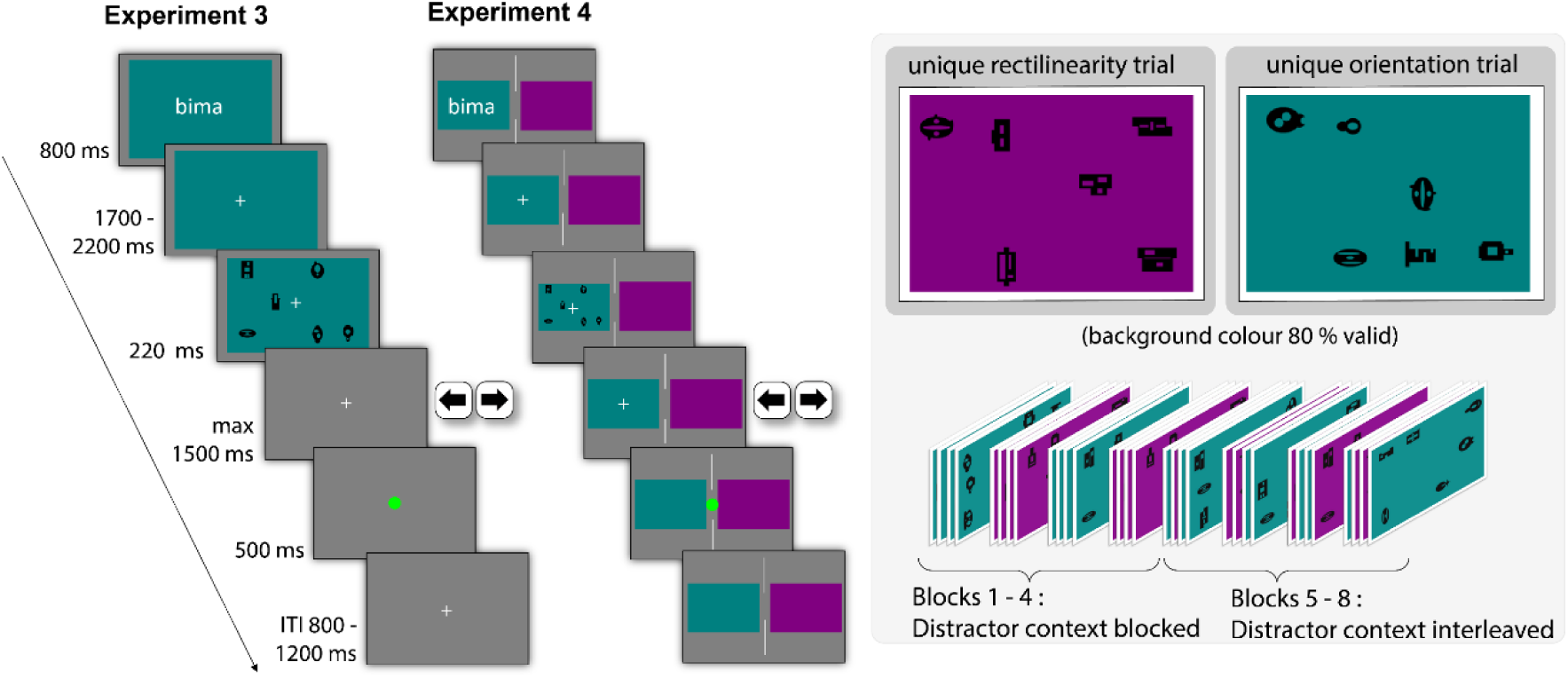
Timeline of a Trial and Design Overview for Experiments 3 and 4 Note. The overall search task was the same as in Experiment 1, but the background colour predicted the diagnostic target dimension with 80% validity. Distractor context was blocked for the first half of the experiment and interleaved for the remaining half. In Experiment 3, background colour changed at the onset of a trial. In Experiment 4, both contexts remained constantly on screen, on the left and right of the display.

The search task consisted of 8 blocks of 40 trials each. To facilitate learning, in the first four blocks, background colour within a block was constant (alternating between blocks), indicating the upcoming trial type with 80% validity. In the second half of the experiment, background colour changed on a trial-by-trial basis, again indicating the upcoming trial type with 80% validity. This design should minimize interference between the two contexts during learning, allowing to establish separate representations and learning rules for both (Flesch et al., 2018). Background colour changed as soon as the name cue was presented on screen and remained visible until the search display had disappeared. Reminders of the associations between distractor contexts and background colour were also shown at the beginning of every block. The experiment took around 50 minutes in total.

Target orientation, rectilinearity and validity were counterbalanced per block, while the specific target individuals and the display side on which the target appeared were counterbalanced across the whole experiment. All factors were counterbalanced separately for both contexts. Which block context participants started with and the associations between block context and background colour were randomized.

#### Stimuli & Setup

Stimuli were created as in Experiment 1, with the exception that the background colour was changed to either blueish green (RGB 0, 127.5, 127.5) or magenta (RGB 127.5, 0, 127.5). The experiment ran online, was programmed in Psychopy and hosted on Pavlovia (https://pavlovia.org). Participants used their own devices to take part in it.

#### Data Analysis

We analysed distractor context effects separately for the blocked and interleaved context blocks with two 2 (trial type: rectilinearity, unique orientation trial) x 2 (context match: matching, mismatching context cue) repeated measures ANOVAs.

To understand how context facilitation evolved within blocks, we compared context facilitation across different trial bins in the blocked context pooled data from all 6-shape trials (all 6 blocks from Experiment 1 and the first 4 blocks of Experiment 3. We computed context facilitation by computing the performance difference between matching and mismatching context trials in each time bin, independent of dimension, and analysed it in a 1 x 4 ANOVA (block time bin: trials 9-16, trials 17-24, trials 25 – 32, trials 32 - 40).

In addition, we tested the effects of intertrial priming on performance in the interleaved condition. For this, we computed performance measures separately for trials in which the diagnostic dimension was the same as on trial n-1 and trials for which the diagnostic dimension had switched from the previous trial, only including those trials on which the previous response had been correct, but independent of the context cues.

Context matching effects for individual dimensions were followed up with t-tests (2-sided, α = .05). Bayes factors for simple t-contrasts and the main effects/interactions were calculated as Bayesian one-sample t-tests for the respective difference scores between conditions or condition averages using the JASP default settings (Cauchy prior with scale 0.707). For comparisons between experiments, Bayesian independent sample t-tests were computed instead.

### Results

Following our preregistration, we excluded trials in which reaction times were below 150 ms or +/- 3 std away from this participant’s mean correct RT. This resulted in a rejection of 1.57% (sd 0.73) of trials.

Overall accuracy was 87.60% (sd 8.18) and mean RT was 702.74 ms (sd 82.40).

Replicating our previous experiments, context expectations facilitated performance when context was blocked (Figure 8; context match: *F*(1,79) = 14.74, *p* < .001, *η_p_²* = 0.157, *BF_10_* = 78.04). Context facilitation depended on which dimension was diagnostic (interaction context match and trial type: *F*(1,79) = 10.95, *p* = 0.001, *η_p_²* = 0.12, *BF_10_* = 16.95), matching context improved performance on unique rectilinearity unique trials (*t*(79) = 5.08, *p* < .001, *d_z_* = 0.57, *BF_10_* > 5000) but not unique orientation trials (*t*(79) = - 0.74, *p* = .46, *d_z_* = -0.08, *BF_10_* = 0.16).

**Figure 8.**
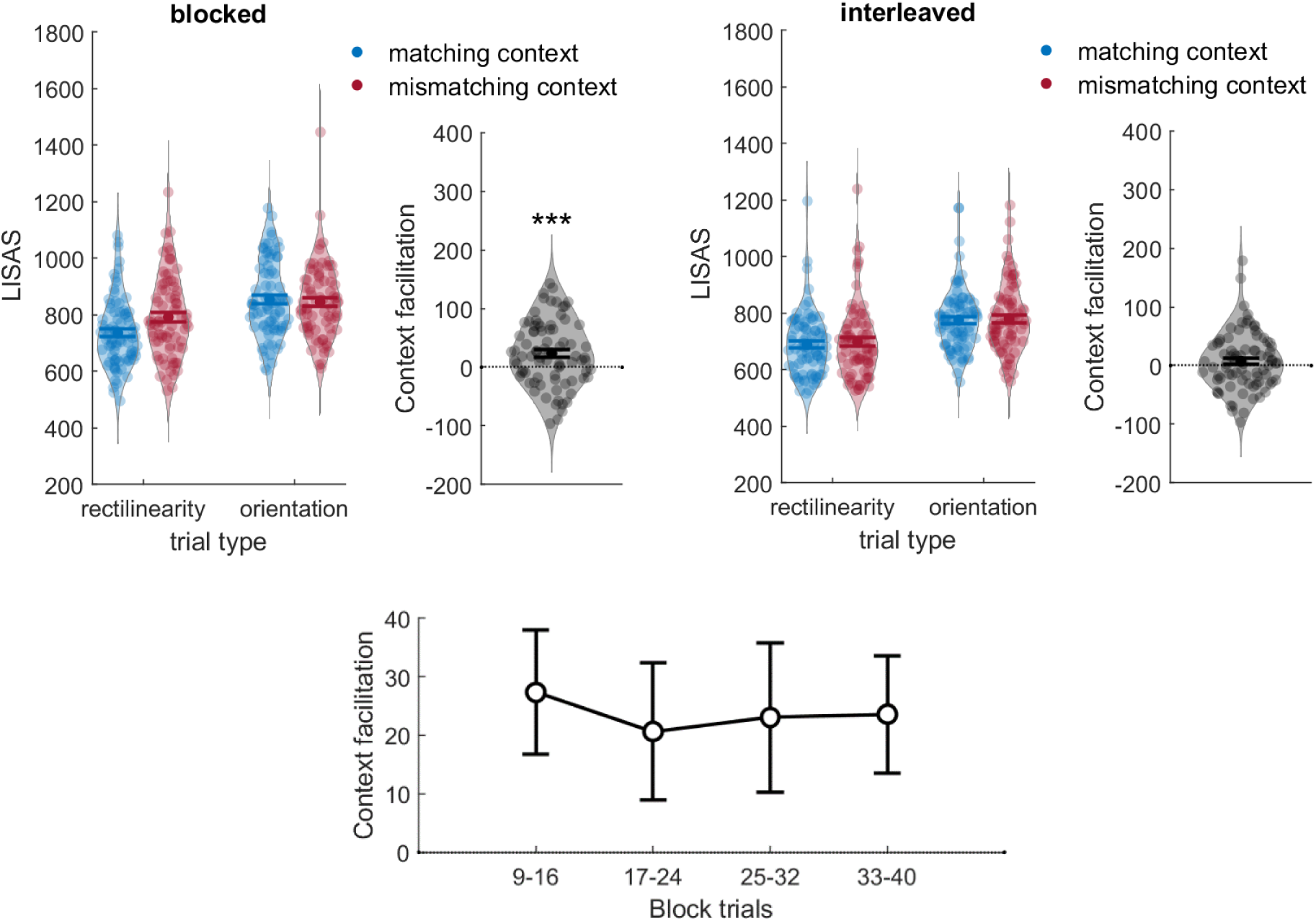
Results of Experiment 3 Note. The upper panel shows performance on both trial types as a function of context (left subpanel) and context facilitation (LISAS mismatching – matching context) averaged across trial types (right subpanel). All error bars are SEM. The lower panel shows context facilitation calculated for individual trial bins of the blocked condition in Experiment 1 and 3 combined.

In the interleaved condition however, where context changed on a trial-by-trial basis, context cues did not influence search performance (main effect context match: *F*(1,79) = 1.68, *p* = 0.20, *η_p_²* = 0.021, *BF_10_* = 0.28), and this was equally the case for both diagnostic dimension (no interaction between context match and trial type: *F*(1,79) = 0.12, *p* = .72, *η_p_²* = 0.00, *BF_10_* = 0.13).

Our results suggested that context may need to be stable over trials in order to influence search performance, leading to two additional exploratory analyses. First, one may ask whether these context expectations could be established relatively quickly within the blocked conditions or gradually developed after multiple trials. We analysed context facilitation across 4 block time bins across pooled data from Experiment 1 and 3, excluding the first 8 trials which were always valid.

Generally, context facilitation remained stable across trial bins (main effect of trial bin: *F*(3,336) = 0.01, *p* = .99, *η_p_²* < 0.001, *BF_10_* = 0.009). In addition, context facilitation already arose in the earliest analysable time bin (trials 9-16, *t*(113) = 2.57, *p* = .01, *d_z_ =* 0.24, *BF_10_* = 2.38), suggesting that participants rapidly adjusted to new distractor contexts.

Second, we analysed whether repeating diagnostic dimensions across trials facilitated performance in an exploratory analysis for the interleaved condition. We computed the LISAS separately for trials in which the diagnostic dimension was the same as on trial n-1 and trials for which the diagnostic dimension had switched from the previous trial, only including those trials on which the previous response had been correct, but independent of the context cues. Indeed, performance tended to be slightly higher when the diagnostic dimension was repeated, although this was not significant (*t*(79) = 1.89, *p* = .06, *d_z_* = 0.21, *BF_10_* = 0.66).

To test whether some participants may still have successfully used the context cues, we also computed the correlation of context facilitation in the blocked and interleaved conditions across participants. This correlation was not significant (*ρ* = - .07, *p* = .52).

Besides the effects of context expectations (or their absence), performance was again always better on unique rectilinearity trials in both blocked and interleaved condition (main effects of trial type, *p*’s < .001).

### Discussion

As in our previous experiments, we found that blocked distractor context modulated the attentional template, with better performance when the diagnostic dimension on a trial was expected rather than unexpected.

When distractor context was interleaved, we no longer found an effect of distractor expectations. While the attentional template was initially biased to emphasize diagnostic target dimensions in the blocked context, it was not influenced by context cues in the interleaved condition and therefore likely included both features of the shapes. Importantly, this was the case even though participants were explicitly informed about the contingencies, were first exposed to them in a blocked manner and reminded about them prior to every block.

Our results showed that explicit context cues were not readily exploited to solve the task, compared to the apparent ease with which regularities in target-distractor regularities were extracted and used when context was blocked. On the one hand, this could suggest that target- distractor relations are generally not learnt in a context-dependent manner. Distractor context effects would then reflect biases based on recent selection history, where target-distractor statistics of previous trials were integrated in the search template independent of context. Accruing over multiple trials, these biases may be stronger than observed in the interleaved condition.

On the other hand, it may still be possible to update the attentional template on a trial-by-trial basis under different conditions. Quickly adjusting to abstract, explicit context cues may be challenging, and not always done by participants, depending perhaps on the amount of training with those cues and whether they are necessary to do the task. In the orientation discrimination study mentioned earlier, participants underwent three experimental sessions (Scolari et al., 2012) and the task required very fine-grained discriminations. The same cues did not affect sensory gain when discrimination was easy (Scolari & Serences, 2009). In a similar vein, when participants searched for a specific watch among multiple instances of another watch, cueing the identity of the distractor watch in advance facilitated search (relative to when there was no cue), but only when participants had not been well familiarized with the target watch prior to search, which may have rendered the task easier (Bravo & Farid, 2016).

While the background colours used were salient, attending to them was not strictly necessary to find the target and they may therefore have been ignored. On the other hand, our results also suggested that context only needed to remain stable over relatively few trials for participants to adjust to context. To test whether it was still possible to emphasize different target dimensions on a trial-by-trial basis, we decided to provide stronger context cues, while keeping the overall task and difficulty equal.

## Experiment 4 – Adjusting to Distractor Context on a Trial-by-Trial Basis with Location as Context Cue

In naturalistic search, different target-distractor relations are often found in different locations, and in addition to visual differences between contexts, spatial location is likely a very strong cue to separate contexts. Memory can strongly depend on spatial context (Godden & Baddeley, 1975; Smith et al., 1978; Smith & Vela, 2001), and spatial separation may therefore help to separate memories of previous searches in both contexts, which may otherwise interfere with each other. If that is the case, combining the different colour contexts with additional (redundant) spatial information, presenting them at different screen locations, may yield stronger context-dependent learning.

Therefore, in Experiment 4 we followed the same approach as in Experiment 3, again testing whether diagnostic target dimensions could be learnt in a context-dependent manner, but distractor contexts were also separated in space. Two coloured squares were continuously presented on the left and right of the screen, with participants’ attention being cued towards one of the two contexts at the start of each trial.

### Methods

#### Participants

80 participants (47 females, mean age: 28.48, sd: 4.23) were included in this new experiment. Additional participants were replaced due to not completing the experiment/having a response rate < 85% (N=4), because their overall performance was < 55% (N=7) or were at chance when a familiar target shape was used as distractor (N=4). Sample size was chosen for consistency with Experiment 3 and again allowed us to detect main effect of context match effect about half as large as the blocked condition in Experiment 1 with 80% power (*η_p_²* > .095, with α = .05).

All participants were recruited via Prolific (https://www.prolific.co), provided informed consent and received £5.00 for their participation. The study was approved by the Radboud University Faculty of Social Sciences Ethics Committee (ECSW-LT-2022-9-9-25489). All data were collected in 2022.

#### Procedure

The overall training and search task were similar to Experiment 3, including the blocked context blocks and the number of trials, with the exception that the two distractor contexts were separated in space (Figure 7). Two coloured squares (blue-green and magenta) were constantly presented on the left and right of the display, separated by a thin white line in the display centre. At the onset of a trial, the name of a shape appeared in one of the squares, designed to direct participants’ fixation towards one context, followed by the search scene. Feedback was given by a coloured fixation dot appearing at the centre of the screen after each trial. The search display always appeared in the cued colour square (while the other one remained empty), but the redundant spatial and colour cue predicted the upcoming trial type with 80% validity.

Target orientation, rectilinearity and validity were counterbalanced per block, while the specific target individuals and the display side on which the target appeared were counterbalanced across the whole experiment. All factors were counterbalanced separately for both contexts. The distractor context participants started with, the associations between block context, background colour and display sides were all randomized.

#### Stimuli & Setup

Search displays were created as in Experiment 3, but their size on screen was reduced (to approximately 15 x 0.9 dva assuming a 57 cm viewing distance). The online experiment was programmed in Psychopy and hosted on Pavlovia (https://pavlovia.org).

#### Data Analysis

Distractor context effects were again analysed separately for the blocked and interleaved context blocks with two 2 (trial type: rectilinearity, unique orientation trial) x 2 (context match: matching, mismatching context cue) repeated measures ANOVAs on the LISAS.

In addition, we compared the interleaved conditions of Experiment 3 and 4, by using a 2 (trial type: rectilinearity, unique orientation trial) x 2 (context match: matching, mismatching context cue) x 2 (Experiment) repeated measures ANOVA, with experiment as between-subjects factor.

Context matching effects for individual dimensions were followed up with t-tests (2-sided, α = .05). Bayes factors for simple t-contrasts and the main effects/interactions were calculated as Bayesian one-sample t-tests for the respective difference scores between conditions or condition averages using the JASP default settings (Cauchy prior with scale 0.707). For comparisons between experiments or groups, Bayesian independent sample t-tests were computed instead.

### Results

Overall accuracy and RT were 86.32% (sd 8.52) and 749.73 ms (sd 114.41), respectively. Following our preregistration and the procedure described for Experiment 3, anticipatory and delayed responses (on average 1.42% (sd 0.86) of trials) were excluded.

Again replicating our previous findings, context expectations influenced search in the blocked condition, with better performance when the diagnostic target dimensions matched participants’ expectations (Figure 9; main effect of context match: *F*(1,79) = 13.32, *p* <.001, *η_p_²* = .14, *BF_10_* = 47.98). This was driven mostly by the unique rectilinearity trials (context match x trial type: *F*(1,79) = 5.04, *p* = .03, *BF_10_* = 1.31; rectilinearity trials: *t*(79) = 3.73, *p* < .001, *d_z_* = 0.417, *BF_10_* = 61.18; orientation trials: *t*(79) = 0.35, *p* = .725, *d_z_* = 0.04, *BF_10_* = 0.13).

**Figure 9.**
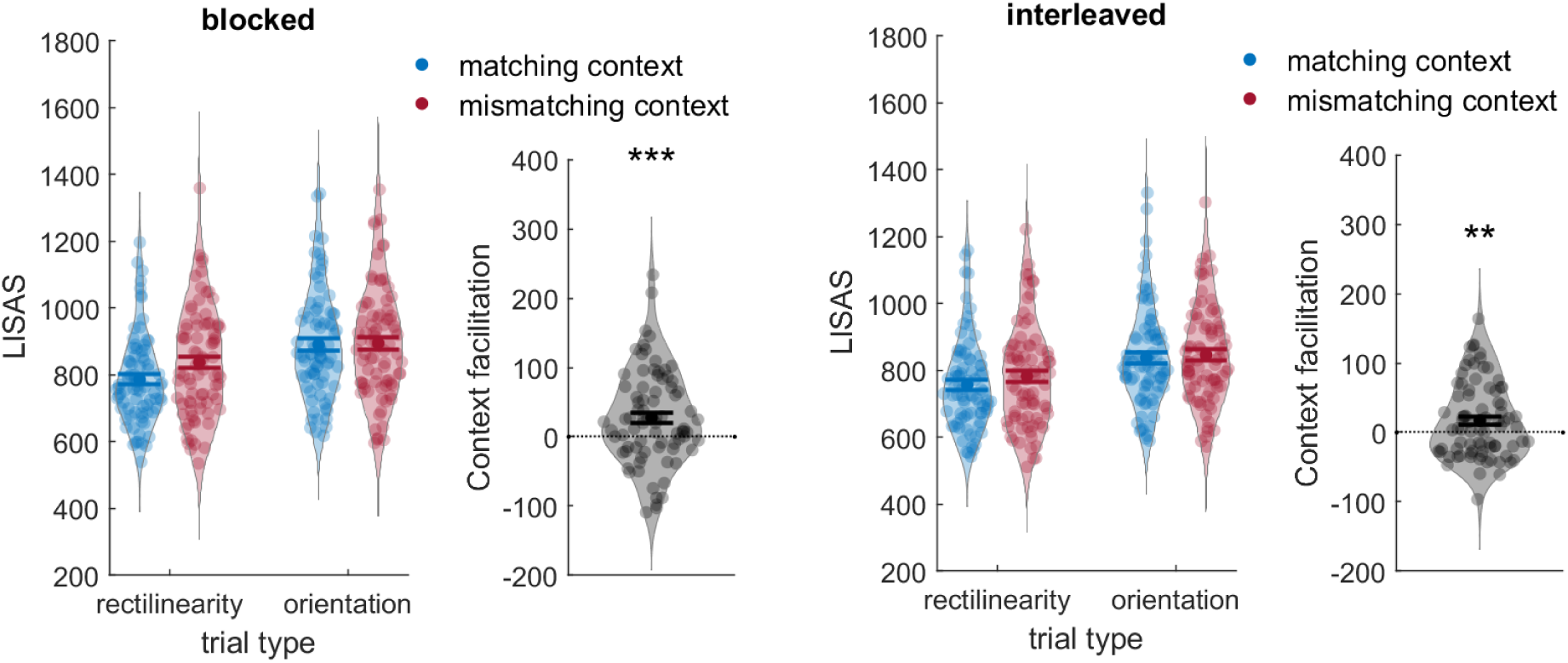
Results of Experiment 4 Note. Performance on both trial types as a function of context (left subpanel) and context facilitation (LISAS mismatching – matching context) averaged across trial types (right subpanel). All error bars are SEM.

Most importantly, the attentional template was also shaped by context cues in the interleaved condition, resulting in better performance when the diagnostic target dimension was expected (context match: *F*(1,79) = 7.98, *p* = .006, *η_p_²* = .09, *BF_10_* = 4.89), with no difference between diagnostic dimensions (context match x trial type: *F*(1,79) = 2.31, *p* = .13, *η_p_²* = .02, *BF_10_* = 0.37; context match unique rectilinearity trials: *t*(79) = 2.90, p = 0.005, *d_z_* = 0.32, *BF_10_* = 5.60; orientation *t*(79) = 1.02, p = 0.311, *d_z_* = 0.11, *BF_10_* = 0.20). Independent of context expectations, the target was always more easily found on rectilinearity unique trials in both the blocked and interleaved condition (p’s < .001).

The effect of context expectations in the interleaved condition remained consistent when combining all data from Experiments 3 and 4 (context match: *F*(1,158) = 8.80, *p* = .003, *η_p_²* = .05, *BF_10_* = 5.85), and the effect did not significantly differ across experiments (context match x experiment: *F*(1,158) = 0.22, *η_p_²* = .01, *BF_10_* = 0.34).

Given that participants formed context expectations in both the blocked and interleaved condition, we again tested whether context facilitation in both conditions correlated across participants, to complement our main preregistered analysis. This was however not significant, neither in Experiment 4 alone (*ρ* = 0.16, *p* = 0.19), nor in Experiments 3 and 4 combined (*ρ* = .06, *p* =.47).

### Discussion

As in Experiment 3, we tested whether target-distractor relations could be learnt in a context-dependent manner and bias the attentional template on a trial-by-trial basis, but using spatial location as additional context cue. Spatial location may support context-depending learning, although a direct comparison between both experiments did not show significantly stronger context facilitation.

Most importantly, we now found that context cues modulated the attentional template on a trial-by-trial basis, indicating that the target-distractor relations were learnt in a contextualized manner and that the attentional template was biased by long term memory for the target-distractor relations in different contexts. Thus, distractor context does not only modulate the attentional template based on context-independent short-term priming mechanisms (including those that may extend over multiple trials), but can also flexibly bias the template, top-down, whenever a familiar distractor context is re-encountered.

Overall, we did not observe strong correlations between context facilitations in blocked and interleaved conditions, even across a large sample (N=160). In stable and rapidly varying distractor contexts, adjusting the attentional template to distractor expectations may not rely on fully overlapping processes. One possibility is that context-independent biases, based on selection history, additionally contribute to context facilitation in the blocked condition but not the interleaved condition.

## General Discussion

Our visual system selects those objects and features for further processing on the basis of their match with an internal representation of our current goals (Duncan & Humphreys, 1989; Eimer, 2014; Wolfe, 2021). Which visual features allow for efficient search, however, depends not only on the visual features of the target, but also on how they relate to the features of the current distractors (Duncan & Humphreys, 1989; Geng & Witkowski, 2019; Navalpakkam & Itti, 2007). To test the flexibility of adaptive biases in attentional templates based on distractor expectations, here we investigated how distractor expectations shape the target template for complex 2D shapes in both blocked and interleaved distractor contexts. Participants searched for these targets in two probabilistic contexts, rendering either the target’s rectilinearity or orientation diagnostic and we tested whether target features in the expected diagnostic dimension were emphasized relative to the non-diagnostic one.

Across multiple experiments, we found that participants learnt the statistical regularities of different distractor contexts when these were blocked, such that search performance was better on trials matching the most frequent diagnostic dimension of a block. The same was found for search trials with only one distractor (Experiment 2), confirming that the attentional template for the target was biased by distractor expectations in a way that emphasized the most diagnostic target dimension. This bias was, however, relative rather than absolute. That is, participants did not simply look for, e.g., any horizontal shape entirely independent of its rectilinearity, as performance on mismatching trials and all 2-shape trials (Experiment 2) was still much higher than chance.

In addition, our results suggest adjusting to distractor context may also be possible without participants noticing these regularities. Over the course of short blocks, participants extracted regularities about common diagnostic feature dimensions from the search scene, independent of specific target or distractor features. Context facilitation was equal for participants who reported being aware of the context manipulation and those who did not (Experiment 2). It is less clear whether the same is true for the interleaved condition in Experiments 3 and 4. The mapping between context cues and target-distractor relations may also be learnable implicitly but we cannot rule out explicit knowledge was necessary in our task. Nonetheless, this knowledge by itself was not sufficient to use context cues in Experiment 3. Further, in our daily life environments, where contexts are highly familiar and rich in cues to separate them, adjusting attentional target templates in a context-dependent may be rather effortless.

Finally, in Experiment 4, (but not Experiment 3), we found that predictive distractor context was also exploited when context changed on a trial-by-trial basis, showing that distractor context also shaped the template independently of biases building up over immediately repeating searches (i.e. selection history). This is relevant as it demonstrates that attentional templates can be truly context-dependent.

Together, these findings provide important evidence that even attentional templates for complex shapes, retrieved from memory, can be flexible to optimally support search performance by highlighting diagnostic information. That being said, it is unlikely that searchers rely on entirely distinct and non-overlapping attentional templates across contexts, as those templates may still be based on the same underlying memory representation (Boettcher et al., 2020; Yu et al., 2023) and within most contexts there will always be some uncertainty about the diagnosticity of certain target features before the object is actually identified. As mentioned already, our results suggest a relative bias in how strongly a given target feature is emphasized in the template, depending on distractor expectations, rather than a total change of the template; accordingly, our effect sizes are relatively small. Whether context cues are used and the template adjusted likely depends on whether context-dependent templates are necessary for successful search (Bravo & Farid, 2016; Scolari et al., 2012; Yu et al., 2023) and how easily separate target representations in different contexts are formed and retrieved from memory (where space could act as a relevant additional context cue). Indeed, the null result we observed for the interleaved condition in Experiment 3 suggests there are also clear limits to the flexibility of the template.

Importantly however, we found that attentional templates were surprisingly flexible under conditions that could have promoted unbiased templates instead, for several reasons. First, targets and distractors were well distinguishable and participants were familiarized with the target shapes outside of a search context, which may promote a distractor-independent representation (cf. Bravo & Farid, 2016). Second, the two dimensions together formed the outline shape of the targets, such that the dimensions were not clearly separable. Third, the use of symbolic cues prevented the selective encoding of one dimension of the current visual input over another (Boettcher et al., 2020), as could be the case when using picture cues. Finally, attentional templates were biased by distractor expectations even when distractor context changed frequently, on a trial-by-trial basis. This indicates that biases in the template can indeed arise relatively efficiently and effortlessly, making it likely that similar mechanisms operate in daily-life visual search.

These findings were obtained from a larger sample of online participants, outside of a typical lab context and are supported by prior studies using similar search tasks with simpler stimuli and a different combination of dimensions (Boettcher et al., 2020; Lee & Geng, 2020). While this provides evidence that context expectations can influence different dimensions of the target template, including rectilinearity as an important dimension for real-world shapes, further dimensions (e.g., size or ‘spikyness’) remain to be tested. Furthermore, it is still unclear whether context facilitation is more strongly observed for target dimensions that are overall prioritized in the template, as suggested by our results in Exp 2A and 3. Indeed, we found differences in context facilitation between target dimensions in some experiments, but these were not consistently present and the two dimensions were not carefully matched in how well they generally allowed to distinguish targets from distractors; it is well possible that orientation is more strongly modulated by context expectations in other settings. More generally, whether distractor expectations bias the template may depend on the relative advantage of including these expectations compared to a stable template (based on the predictability of specific distractor and target features within and across different contexts). Difficult search tasks with few, highly predictable distractor contexts and dimensions that can be independently represented by the visual system may lead to stronger context effects than searches across many variable and less predictable contexts.

We interpret our results in terms of biased attentional templates rather than, for example, improvements in distractor grouping (Experiment 2). However, independent of grouping, in principle it is possible that expected distractor features were suppressed better (see van Moorselaar & Slagter, 2020 for a review) rather than target features being enhanced. We think this is unlikely, because cueing distractor features on a trial-by-trial basis often leads to no benefits, or even costs (e.g., M. W. Becker et al., 2016; Cunningham & Egeth, 2016). Furthermore, while feature-based inhibition based on statistical learning and blocked distractors is possible (Stilwell et al., 2019; van Moorselaar et al., 2021; Van Moorselaar et al., 2020; Vatterott & Vecera, 2012), it is likely not context-dependent (see Britton & Anderson, 2019; de Waard et al., 2022 for space-based suppression). Successful feature-based distractor inhibition in our experiments would require learning specific distractor features associated with every target in both contexts. Enhancement of diagnostic target features therefore seems a more probable mechanism.

Where and when does this enhancement take place? A relevant direction for future research will be to establish whether distractor context expectations influence early attentional selection and guidance, or alternatively only later decision processes. Attentional selection and target-identification might rely on different templates (Wolfe, 2021) and those may be differentially shaped by distractor context (Hamblin-Frohman & Becker, 2021; Yu et al., 2022). At the neural level, attentional templates have been associated with a baseline increase of neural activity in neurons tuned to target features (Desimone & Duncan, 1995), which serves to bias competition in favour of the target in a subsequently presented search array (see e.g. Battistoni et al., 2017; Eimer, 2014 for reviews). A bias in the template could thus be instantiated by an increase in preparatory activity of neurons coding for the discriminative feature of the target and lead to an increased response of these neurons when the target is presented. In line with this, Scolari et al. (2012) found that the gain of V1 neurons whose activity discriminated best between target and distractors in a difficult orientation discrimination task was selectively enhanced. In the current study, looking for, e.g., a rectilinear shape in rectilinearity unique trials could have increased baseline activity or gain of neurons preferring rectilinear features. Context-dependent preparatory biases may influence early sensory processes, but additional neuroimaging studies are needed to understand where in the visual hierarchy and at which processing stages context expectations influence search.

During real-world search, the attentional template may be informed by context also in other ways. For example, when experiencing the same object (or object category) across multiple contexts, we may learn which features are diagnostic of this object across different contexts and selectively include those diagnostic parts in the template (Reeder & Peelen, 2013). Which features are diagnostic may not only reflect the relation of targets and distractors, but also the variability of the target itself (i.e., which features consistently characterize exemplars of a category or the same object across time). Along these lines, there is evidence that target-dimensions with lower variance are emphasized in the template for target-match decisions (Witkowski & Geng, 2019, 2022). This can be seen as long-term biases based on extensive experience. While there may therefore be less need to adjust these object representations to respective contexts, they are still not static and can also be biased by current context and expectations. Predictive distractor contexts are only one instance of regularities shaping the attentional template. For example, recent studies found that participants are sensitive to colour statistics within specific object categories (Bahle et al., 2021) also in a context-dependent manner (Kershner & Hollingworth, 2022). In addition, relations between objects and the overall scene context (e.g., the expected retinal size of an object at a particular location) are encoded in the attentional template (Gayet & Peelen, 2022), all providing evidence for the adaptability of attentional templates.

To conclude, we provide evidence for flexible and adaptive biases in the attentional template when searching for the same objects within different contexts. Distractor context was learnt quickly and seemed to be incorporated in the template independently of awareness of the context manipulation. Moreover, attentional templates were biased by distractor expectations in a context-dependent manner, even when context was interleaved. Further investigating how and when context is used to shape attentional allocation will help to understand how we efficiently find objects in our daily-life environments.

## Appendix

### Results Accuracy & RT – Experiment 1

We conducted the same 2 (trial type: unique rectilinearity, unique orientation trial) x 2 (context match: presented in matching, mismatching context block) repeated-measures ANOVA reported for the LISAS in the main text also separately for accuracy and RT.

For accuracy, the main effect of context match was significant (*F*(1,33) = 8.05, *p* = .008, *η_p_²* = .20, *BF_10_* = 5.37). Averaged over trial types, context facilitation was 3.54% (sd 7.16). In addition, there was a significant main effect of trial type, with better performance on rectilinearity unique trials (*F*(1,33) = 8.53, *p* = .006, *η_p_²* = .21, *BF_10_* = 6.45). Both factors did not interact (*F*(1,33) = 0.2, *p* = .65, *η_p_²* = .01, *BF_10_* = 0.20; context match unique rectilinearity trials alone (*t*(33) = 1.68, *p* = .10, d_z_ = 0.29*, BF_10_* = 0.65); unique orientation trials (*t*(33) = 2.32, *p* = .03, *d_z_* = 0.29, *BF_10_* = 1.89).

For reaction times, context facilitation was 7.50 ms (sd 34.64), but the main effect of context match not significant (*F*(1,33) = 1.59, *p* = .22, *η_p_²* = .05, *BF_10_* = 0.38). There was also a significant main effect of trial type (*F*(1,33) = 78.52, *p* < .001, *η_p_²* = .70, *BF_10_* > 3 x 10^7^). There was no significant interaction between trial type and context match (*F*(1,33) = 0.14, *p* = .71, *η_p_²* = .004, *BF_10_* = 0.20; unique rectilinearity: *t*(33) = 1.00, *p* = .34, *d_z_* = 0.17, *BF_10_* = 0.29, unique orientation: *t*(33) = 0.47, *p* = .64, *d_z_* = 0.08, *BF_10_* = 0.20).

### Results Accuracy & RT – Experiment 2

#### Experiment 2A

For accuracy, there was no significant main effect of context match (*F*(1,33) = 0.3, *p* = .59, *η_p_²* = .01, *BF_10_* = 0.21). The main effect of trial type was significant (*F*(1,33) = 5.85, *p* = .02, *η_p_²* = .15, *BF_10_* = 2.21), due to better performance on rectilinearity unique trials. Both factors did not interact (*F*(1,33) = 0.09, *p* = .76, *η_p_²* = .00, *BF_10_* = 0.19; context match unique rectilinearity trials: *t*(33) = -0.69, *d_z_* = -0.12, *p* = .50, *BF_10_* = 0.23; orientation trials: t(33) = -0.12, *d_z_* = -0.02, *p* = .91, *BF_10_* = 0.19).

For reaction times, we found a significant main effect of match (*F*(1,33) = 4.65, *p* = .038, *η_p_²* = .12, *BF_10_* = 1.41), reflecting a context facilitation of 10.87 ms (sd 29.36) across trial types. There was also a significant main effect of trial type (*F*(1,33) = 46.43, *p* < .001, *η_p_²* = .58, *BF_10_* > 1×10^7^) and a significant interaction between both (*F*(1,33) = 8.01, *p* = .008, *η_p_²* = .20, *BF_10_* = 5.30). This reflected significant context facilitation for the rectilinearity trials alone (*t*(1,33), 3.61, *p* = .001, *d_z_* = 0.62, *BF_10_* = 32.10) but no facilitation in the orientation trials (*t*(1,33) = -1.26, *p* = .21, *d_z_* = - 0.21, *BF_10_* = 0.38).

#### Experiment 2B

In accuracy, the main effect of context match was not significant (*F*(1,79) = 1.53, *p* = .21, *η_p_²* = .019, *BF_10_* = 0.26). As performance was better on rectilinearity unique trials, the main effect of trial type was significant (*F*(1,79) = 16.77, *p* < .001, *η_p_²* = .175, *BF_10_* = 196.68). There was no significant interaction between both (*F*(1,33) = 1.53, *p* = .21, *η_p_²* = .004, *BF_10_* = 0.15; context match rectilinearity trials: *t*(79) = 0.33, *p* = .74, *d_z_* = 0.03, *BF_10_* = 0.13; orientation trials: *t*(79) = 1.21, *p* = .29, *d_z_* = 0.14, *BF_10_* = 0.25).

For RT, the main effect of context match was significant (*F*(1,79) = 13.38, *p* < .001, *η_p_²* = .145, *BF_10_* = 28.46), reflecting a context facilitation of 10.89 ms (sd 26.61). We also found a significant main effect of trial type (*F*(1,79) = 137.98, *p* < .001, *η_p_²* = .63, *BF_10_* > 5 x 10^15^). Trial type and context match did not interact significantly (*F*(1,79) = 0.11, *p* = .74, *η_p_²* = .00, *BF_10_* = 0.13; context match rectilinearity trials: *t*(79) = 1.60, *p* = .11, *d_z_* = 0.21, *d_z_* = 0.18, *BF_10_* = 0.42; orientation trials *t*(79) = 1.92 *p* = .06, *d_z_* = 0.21, *BF_10_* = 0.71).

Comparing context facilitation for participants who had high or no awareness of the context manipulation combined for all participants of Experiment 2A and 2B, context facilitation for RT was not significant for the no-awareness group (*F*(1,34) = 0.83, *p* = .37, *η_p_²* = .25, *BF_10_* = 0.27), but was found for the high-awareness group (*F*(1,33) = 10.76, *p* = .003, *η_p_²* = .25, *BF_10_* = 14.51) and the difference between both did not reach significance (*t*(67) = -1.93, *p* = .06, *d* = -0.46, *BF_10_* = 1.18).

### Results Accuracy & RT – Experiment 3

In the blocked context blocks, we observed a main effect of context match (*F*(1,79) = 5.69, *η_p_²* = 0.067, *p* = 0.02, *BF_10_* = 1.76) in accuracy, with context facilitation of 2.11% (sd 7.91) averaged over trial types. There was also a main effect of trial type, due to better performance for rectilinearity unique trials (*F*(1,79) = 25.32, *p* < .001, *η_p_²* = .248, *BF_10_* = 147.00). The interaction between trial type and context match was not significant (*F*(1,79) = 1.84, *p* = .18, *η_p_²* = .022, *BF_10_* = 0.30; context match: unique rectilinearity trials: *t*(79) = 2.94, *p* < 0.01; unique orientation: *t*(79) = 0.46, *p* = 0.65; context match unique orientation trials (*t*(79) = 2.94, p = 0.004, *d_z_* = 0.33, *BF_10_ =* 6.57, unique orientation trials t(79) = 0.46, p = 0.647, *d_z_* = 0.05, *BF_10_ =* 0.13).

For RT, there was also a main effect of context match, context (*F*(1,79) = 6.81, *p* = .01, *η_p_²* = .079, *BF_10_* = 2.68) reflecting a 12.60 ms (sd 43.16) context advantage averaged over trial types. There was also a main effect of trial type type (*F*(1,79) = 82.507, *p* < .0001, *η_p_²* = .510, *BF_10_* > 4000), and a significant interaction between both (*F*(1,79) = 6.818, *p* = .001, *η­²* = .123, *BF_10_* = 24.30). This reflected a significant match effect for rectilinearity alone (*t*(1,79) = 4.36, *p* < .0001, *d_z_* = 0.487, *BF_10_* = 438.53), but a numerically negative context facilitation for orientation (*t*(1,79) = - 1.58, *p* = .11, *d_z_* = 0.18, *BF_10_* = 0.41)

In the interleaved condition, the main effect of context match did not reach significance for accuracy (*F*(1,79) = 3.11, *p* = .08, *η_p_²* = .038, *BF_10_* = 0.54), reflecting a 1.17 % (sd 5.91) context advantage. The main effect of trial type was again significant (*F*(1,79) = 29.02, *p* < .001, *η_p_²* = 0.269, *BF_10_* = 86.04) and there was no interaction between both (*F*(1,79) = 2.37, *p* = .12, *η_p_²* = .029, *BF_10_* = 0.38; context match unique rectilinearity: *t*(79) = 0.32, *p* = 0.75, d_z_ = 0.04, *BF_10_* = 0.13; unique orientation: *t*(79) = 2.16, *p* = 0.03, *d_z_* _=_ 0.24, *BF_10_* = 1.11). There was no significant effect of the diagnostic dimension of the previous trial on accuracy in the current trial (*t*(79) = 1.23, *p* = 0.22, *d_z_* = 0.14, *BF_10_* = 0.25).

For RT, the main effect of match was not significant (*F*(1,79) = 0.08, *p* = .76, *η_p_²* = .001, *BF_10_* = 0.13). There was, however, again a main effect of trial type (*F*(1,33) = 107.3, *p* < .001, *η_p_²* = .58, *BF_10_* > 1 x 10^6^) and no significant interaction between both factors (*F*(1,79) = 2.65, *p* = .11, *η_p_²* = .032, *BF_10_* = 0.43; context match unique rectilinearity trials: *t*(79) = 1.10, p = 0.273, *d_z_* = 0.12, *BF_10_* = 0.22 ; unique orientation: *t*(79) = -1.32, *p* = 0.19, *d_z_* = - 0.15, *BF_10_* = 0.28). The diagnostic dimension of the previous trial did also not significantly affect RT on the current trial (*t*(79) = 1.00, *p* = 0.32, *d_z_* = 0.11, *BF_10_* = 0.20). Context facilitation in both conditions did not correlate significantly (accuracy: ρ = .02, *p* = .86; RT: ρ = .03, *p* = 0.78).

### Results Accuracy & RT – Experiment 4

In the blocked condition, the main effect of context match for accuracy was significant (*F*(1,79) = 11.50, p = 0.001, *ηp²* = 0.13, *BF_10_* = 22.43), reflecting an overall advantage of 3.37 % (sd 8.90) for trials in matching contexts. We also found a main effect of trial type (*F*(1,79) = 32.35, *p* < .001, *ηp²* = 0.29, *BF_10_* = 602.43) and no interaction between both (*F*(1,79) =1.11, *p* = .27, *η_p_²* = .01, *BF_10_* = 0.21; context match rectilinearity trials: *t*(79) = 3.09, *p* = 0.003, *d_z_* = 0.35, *BF_10_* = 9.73; orientation*: t*(79) = 1.62, *p* = 0.11, *d_z_* = 0.18, *BF_10_* = 0.43).

For RT the main effect of context match failed to reach significance (*F*(1,79) = 3.53, *p* = .06, *η_p_²* = .04, *BF_10_* = 0.67) and the main effect of trial type again significant (*F*(1,79) = 74.00, *p* < .001, *η_p_²* = .48, *BF_10_* = 19.62). Both factors interacted (*F*(1,79) = 5.27, *p* = .02, *η_p_²* = .06, *BF_10_* = 1.46), reflecting significant context facilitation for unique rectilinearity trials (*t*(79) = 3.08, *p* = .003, *d* = 0.34, BF_10_ = 9.54), but not unique orientation trials (*t*(79) = -0.86, *p* = .39, *d_z_* = -0.10, BF_10_ = 0.18).

In the interleaved condition, we also found a significant effect of context match on accuracy (*F*(1,79) = 7.41, *p* = .008, *η_p_²* = .09, *BF_10_* = 3.81), with overall context facilitation of 2.09% (sd 6.85). There was also a significant main effect of trial type (*F*(1,79) = 28.18, *p* < .001, *ηp²* = .26, *BF_10_* = 37.65), but no interaction between both (*F*(1,79) = 0.48, *p* = .49, *η_p_²* = .01, *BF_10_* = 0.16; context match unique rectilinearity: *t*(79) = 1.51, *p* = 0.13, *d_z_* = 0.24, *BF_10_* = 0.36 ; orientation: *t*(79) = 2.17, p = 0.03, *d_z_* = 0.24, *d_z_* = 0.17, BF_10_ = 1.13).

For RT, the overall effect of context match was not significant (*F*(1,79) = 1.17, *p* = .28, *η_p_²* = .01, *BF_10_* = 0.22). There was, however, again a main effect of trial type (*F*(1,79) = 74.94, *p* < .0001, *η_p_²* = .49, *BF_10_* = 1.02) and a significant interaction between trial type and context match type (*F*(1,79) = 4.49, *p* = .04, *η_p_²* = .05, *BF_10_* = 0.02). Context facilitation was significant for unique rectilinearity trials (*t*(79) = 2.39, *p* = .02, *d_z_* = 0.27, BF_10_ = 1.78), but not unique orientation trials (*t*(79) = -0.88, *p* = .38, *d_z_* = -0.10, BF_10_ = 0.18).

Context facilitation in both conditions did not correlate significantly (accuracy: *ρ* = .18, *p* = 0.18; RT: *ρ* = .14, *p* = 0.21).

## Footnotes

1 While rotation generally preserves shape, here orientation may also be seen as elongation, and our vertically or horizontally oriented shapes classified as either wide or long shapes.

