## Supplementary Figures for "Expected Distractor Context Biases the Attentional Template for Target Shapes"

**Supplementary Material**

**Figure S1**

*Results of Experiment 1*

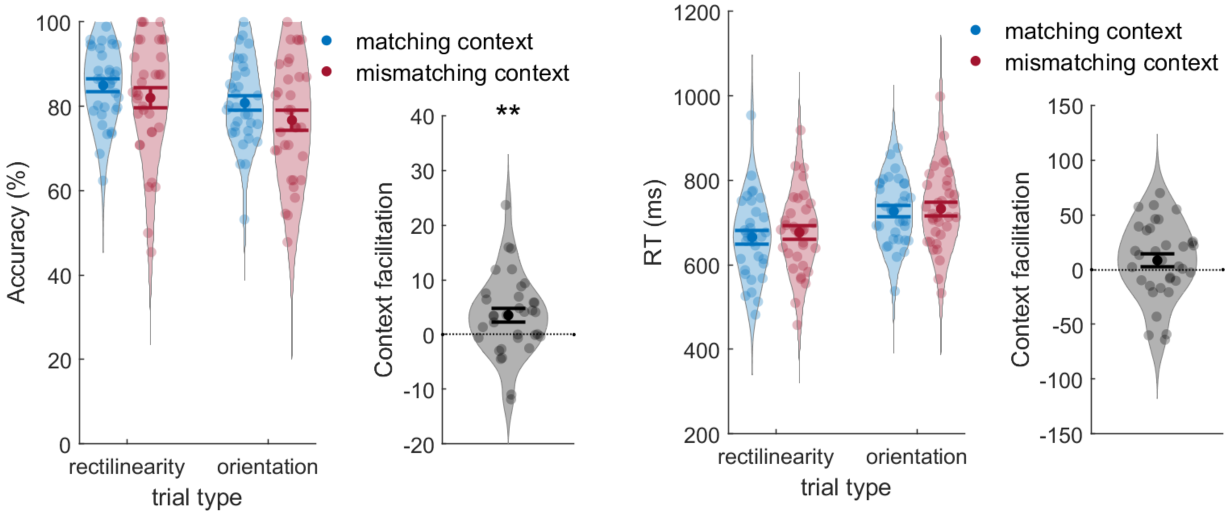


*Note.* Performance (accuracy and RT) on both trial types as a function of block context (left subpanel) and context facilitation (difference between matching and mismatching) averaged across trial types (right). All error bars are SEM.

**Figure S2**

*Results of Experiment 2A & 2B*

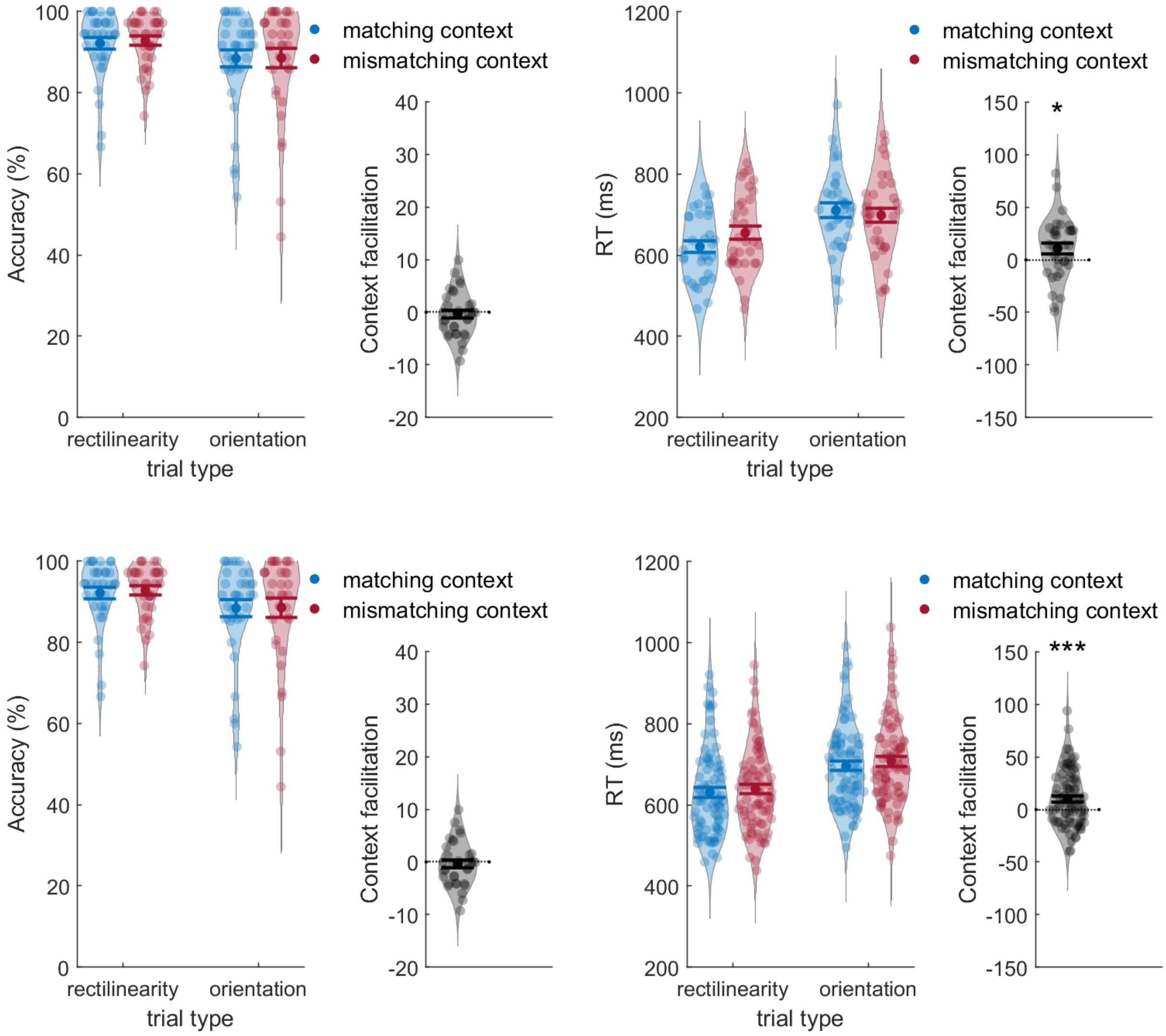


*Note.* Performance (accuracy and RT) on both trial types as a function of block context (left subpanel) and context facilitation (difference between matching and mismatching) averaged across trial types (right). Upper panels show results for Experiment 2A, lower for Experiment 2B, All error bars are SEM.

**Figure S3**

*Results of Experiment 3*

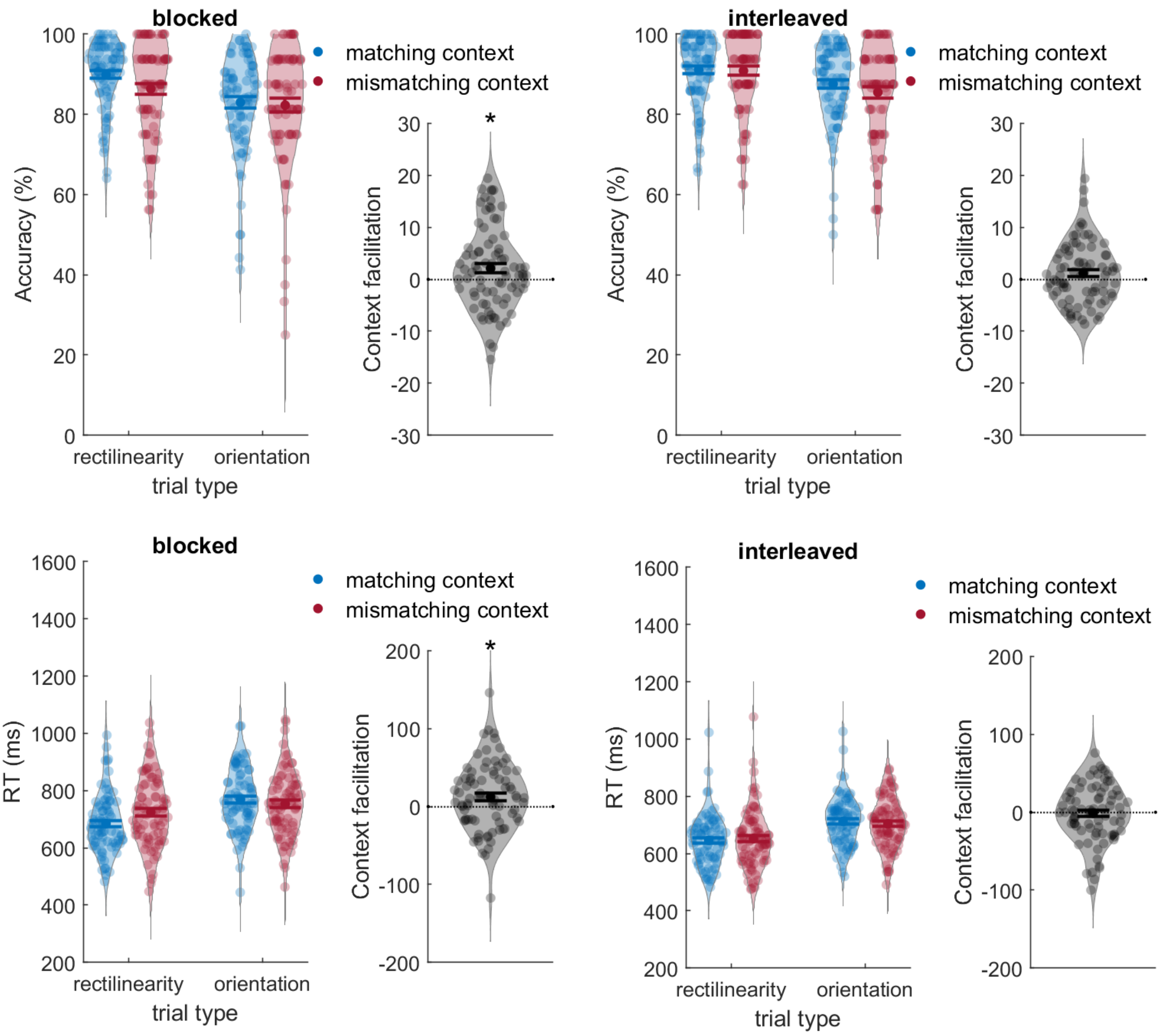


*Note* Performance (accuracy and RT) on both trial types as a function of context (left subpanel) and context facilitation (difference between matching and mismatching contexts) averaged across trial types (right subpanel). All error bars are SEM.

**Figure S4**

*Results of Experiment 4*

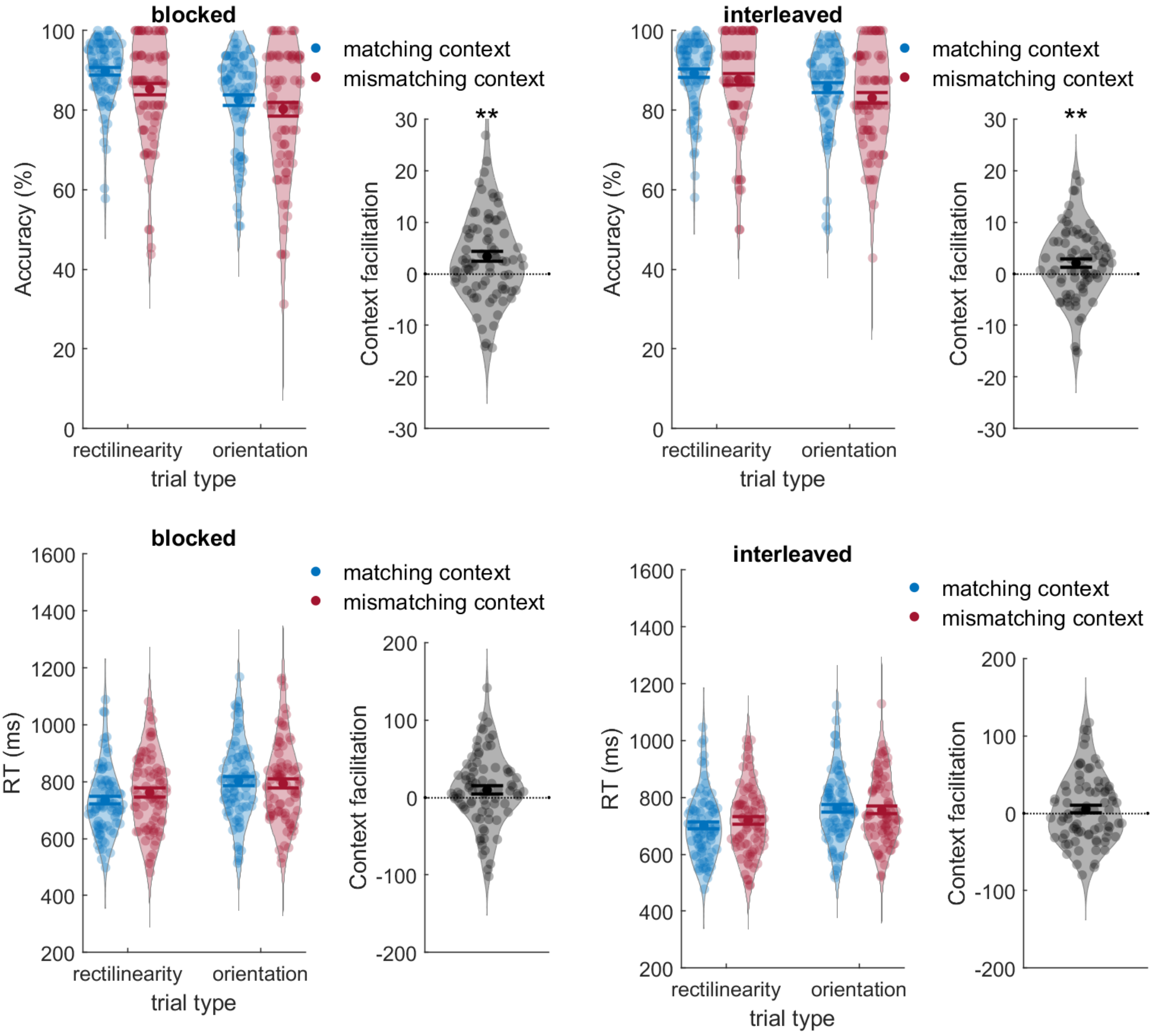


*Note* Performance (accuracy and RT) on both trial types as a function of context (left subpanel) and context facilitation (difference between matching and mismatching contexts) averaged across trial types (right subpanel). All error bars are SEM.
